# Sex-specific lesion topographies explain outcomes after acute ischemic stroke

**DOI:** 10.1101/2020.09.25.308742

**Authors:** Anna K. Bonkhoff, Markus D. Schirmer, Martin Bretzner, Sungmin Hong, Robert W. Regenhardt, Mikael Brudfors, Kathleen L. Donahue, Marco J. Nardin, Adrian V. Dalca, Anne-Katrin Giese, Mark R. Etherton, Brandon L. Hancock, Steven J. T. Mocking, Elissa C. McIntosh, John Attia, Oscar R. Benavente, Stephen Bevan, John W. Cole, Amanda Donatti, Christoph J. Griessenauer, Laura Heitsch, Lukas Holmegaard, Katarina Jood, Jordi Jimenez-Conde, Steven J. Kittner, Robin Lemmens, Christopher R. Levi, Caitrin W. McDonough, James F. Meschia, Chia-Ling Phuah, Arndt Rolfs, Stefan Ropele, Jonathan Rosand, Jaume Roquer, Tatjana Rundek, Ralph L. Sacco, Reinhold Schmidt, Pankaj Sharma, Agnieszka Slowik, Martin Söderholm, Alessandro Sousa, Tara M. Stanne, Daniel Strbian, Turgut Tatlisumak, Vincent Thijs, Achala Vagal, Johan Wasselius, Daniel Woo, Ramin Zand, Patrick F. McArdle, Bradford B. Worrall, Christina Jern, Arne Lindgren, Jane Maguire, Danilo Bzdok, Ona Wu, Natalia S. Rost, on behalf of the MRI-GENIE and GISCOME Investigators and the International Stroke Genetics Consortium

## Abstract

Acute ischemic stroke affects men and women differently in many ways. In particular, women are oftentimes reported to experience a higher acute stroke severity than men. Here, we derived a low-dimensional representation of anatomical stroke lesions and designed a sex-aware Bayesian hierarchical modelling framework for a large-scale, well phenotyped stroke cohort. This framework was tailored to carefully estimate possible sex differences in lesion patterns explaining acute stroke severity (NIHSS) in 1,058 patients (39% female). Anatomical regions known to subserve motor and language functions emerged as relevant regions for both men and women. Female patients, however, presented a more widespread pattern of stroke severity-relevant lesions than male patients. Furthermore, particularly lesions in the posterior circulation of the *left* hemisphere underlay a higher stroke severity exclusively in women. These sex-sensitive lesion pattern effects could be discovered and subsequently robustly replicated in two large independent, multisite lesion datasets. The constellation of findings has several important conceptual and clinical implications: 1) suggesting sex-specific functional cerebral asymmetries, and 2) motivating a sex-stratified approach to management of acute ischemic stroke. To go beyond sex-averaged stroke research, future studies should explicitly test whether acute therapies administered on the basis of sex-specific cutoff volumes of salvageable tissue will lead to improved outcomes in women after acute ischemic stroke.

## Introduction

Stroke affects more than 15 million people each year.^1^ It is known to result in a substantial overall degree of long-term impairment across men and women.^23^ However, numerous epidemiological studies indicate clinically relevant, sex-related differences in the characteristics of ischemic cerebrovascular disease.^4, 5^ For instance, women have a lower stroke incidence than men when younger, yet this initially low women-to-men incidence ratio is decisively reversed in the oldest age groups (>85 years).^6^ This drastic increase in stroke incidence in older women indicates why expected demographical changes, i.e., an aging population, will affect men and women differently: In the US, projections suggest ∼200.000 more disabled women after stroke than men by 2030.^7^

Further sex differences relate to women more often presenting with non-classic stroke symptoms, such as fatigue or changes in mental status^8, 9^ and having a higher risk of delays in hospital arrival.^10, 11^ Also, women feature a higher risk of cardioembolic stroke due to atrial fibrillation,^12^ which may contribute to the often observed higher acute ischemic stroke severity in female patients.^13^ However, this excess in stroke severity in women has been found to persist even after adjusting for their greater age at onset.^14^ Importantly, women seem to experience more serious strokes despite comparable lesion sizes in men and women.^15^ In fact, a similar observation of sex-specific lesion volume effects was noted in case of aphasia, as smaller lesion volumes were sufficient to cause aphasia in women, while larger ones were required in men.^16^

Going beyond lesion volume, lesion symptom mapping studies have enriched our understanding of anatomically unique lesion locations underlying specific symptoms post-stroke.^17, 18, 19^ In case of stroke severity, these analyses have determined widespread lesions in white matter, basal ganglia, pre- and postcentral gyri, opercular, insular and inferior frontal regions to be most relevant for a higher stroke severity, especially if affecting the left hemisphere.^20^ While these lesion symptom studies have enabled to uncover eloquent lesion locations in high spatial resolution, they have been systematically blind to any potential sex disparities. If considered at all, sex was treated as a nuisance source of no interest and regressed out prior to the main analysis.^20^ Thus, none of the recently employed analytical approaches in clinical neuroimaging allowed for a dedicated, explicit investigation of sex-specific lesion pattern effects in relation to continuous outcome scores.

To address these previous methodological constraints, the aim of the current study was to design and conduct a lesion symptom analysis that was capable of capturing male and female-specific lesion patterns underlying stroke severity in a statistically robust and spatially precise manner. For this purpose, we leveraged neuroimaging data originating from two large, independent hospital-based cohorts gathering data of 1,058 acute ischemic stroke (AIS) patients. We tailored and deployed sex-aware hierarchical Bayesian models to simulate predictions of acute ischemic stroke severity and to elucidate the sex-specific effects of lesion patterns affecting similar brain regions in women and men. We hypothesized to derive lesion constellations underpinning more severe stroke especially in women, potentially indicating sex-specific brain function maps on the one hand and facilitating more sex-aware acute stroke treatment decisions on the other hand. Such a sex-informed acute stroke care could eventually alleviate the burden of disease on an individual patient level, yet also broader and socioeconomically relevant level.

## Results

We here present a generative analysis of acute stroke severity, putting a particular focus on sex-specific lesion pattern effects. As in previous work,^21^ we successively combined 1) the automated low-dimensional embedding of high-dimensional lesion information via non-negative matrix factorization (NMF)^22^, and 2) probabilistic modeling to simulate the prediction of the acute stroke severity, as measured by the National Institute of Health Stroke Scale (NIHSS). We thus first determined pivotal, general lesion pattern effects across all subjects and successively concentrated on similarities and differences between men and women. We interpret explanatory relevances on the level of NMF-derived low dimensional lesion representations, that we call *lesion atoms*, as well as the same relevances transformed back to the level of the anatomical grey matter brain regions and white matter tracts.

### Stroke sample characteristics

The derivation cohort considered 208 female and 347 male patients (n=555 in total, 37% females). The main outcome of interest was the acute NIHSS-based stroke severity within the first 48 hours after stroke onset (mean(SD): 5.03 (5.95)). More than one fourth of stroke subjects had a recorded hypertension (28.1%), 19.5% had the diagnosis of diabetes mellitus, 6.3% atrial fibrillation and 7.6% a pre-stroke diagnosed coronary artery disease. As expected based on prior reports,^12^ women had a higher absolute frequency of atrial fibrillation than men (9.1% vs. 4.6%; c.f. **Table 1** for further sex-specific numbers). Acute stroke lesion volume did not differ significantly between men and women (two-sided t-test: *p*=0.98). Moreover, the numbers of lesioned voxels within each cortical or subcortical grey matter brain region or white matter tract did neither differ significantly between the sex categories, nor between the left and right hemisphere (all two-sided t-tests *p*>0.05, Bonferroni-corrected for multiple comparisons, c.f. **Supplementary Tables 1 and 2**).

**Table 1.**
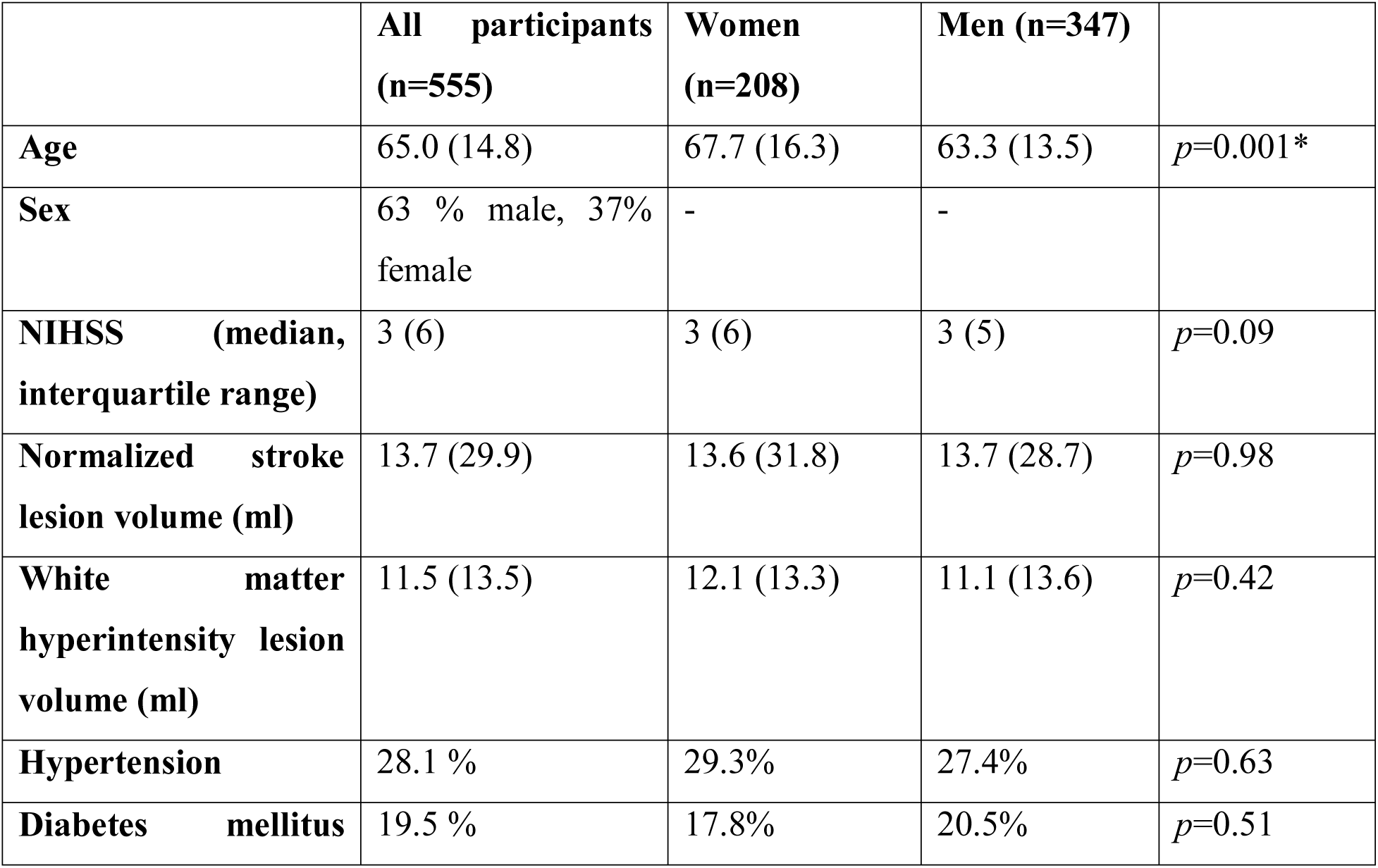
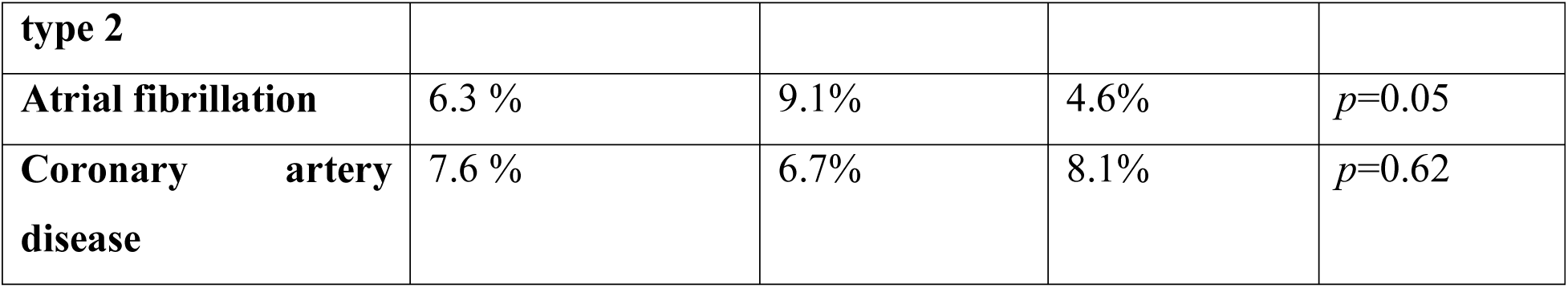
Patient characteristics. Mean (SD) unless otherwise noted. The groups of male and female patients were compared via two sample t-tests or fisher’s exact test as appropriate. **Asterisks indicate significant differences between men and women after Bonferroni-correction for eight multiple comparisons.**

### Anatomy of the extracted lesion atoms in stroke patient

We reduced the high-dimensional lesion voxel-space by first computing the number of lesioned voxels within each of 109 cortical and subcortical grey matter regions, as well as 20 white matter tracts and subsequently employing unsupervised non-negative matrix factorization to ten final *lesion atoms*. Derived low-dimensional *lesion atoms* were found to represent anatomically plausible, hemisphere-specific lesion patterns. The centers of these lesion patterns varied from anterior to posterior and subcortical to cortical regions and were broadly similar between hemispheres (**Figure 1A & B**). In correspondence to the primary distribution of individual lesions, most of these *lesion atoms* related to infarcts in the left and right MCA-supply territories and to a lesser extent to infarcts in the posterior circulation. The maximum lesion overlap was localized in left and right subcortical MCA and insular region (**Figure 1C**). Subcortical and cortical lesion patterns were represented in separate *lesion atoms* in the right hemisphere, while they were captured in a joint lesion atom on the left. Therefore, we obtained six *lesion atoms* in the right and four *atoms* in the left hemisphere.

**Figure 1.**
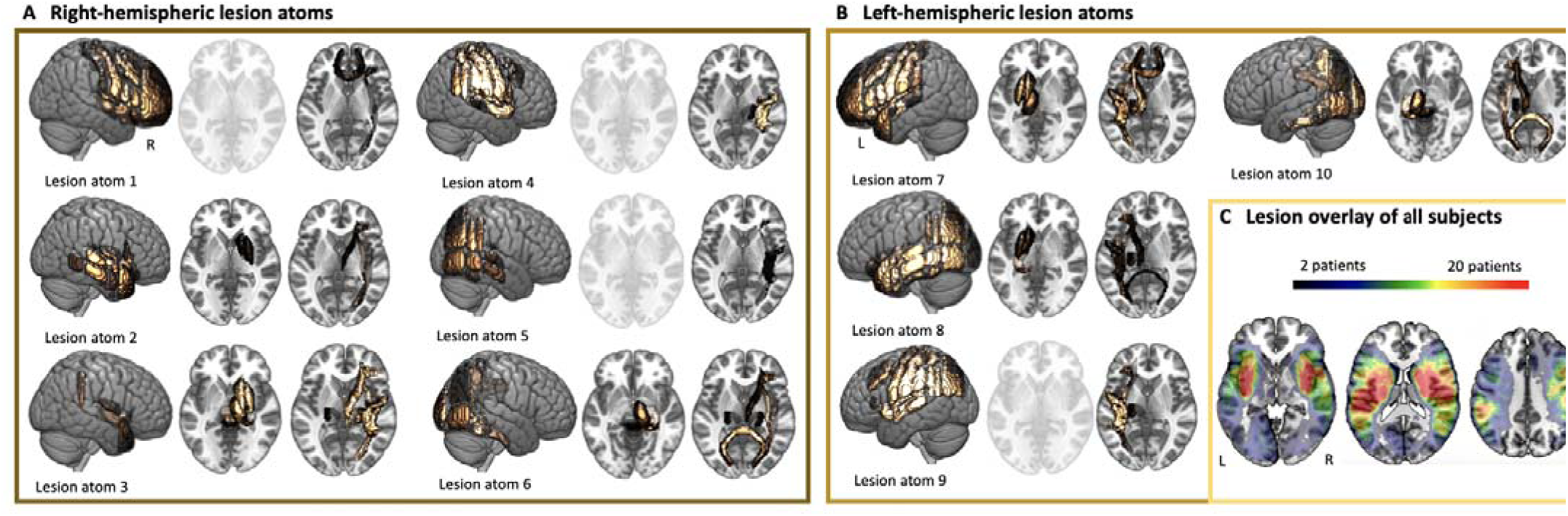
Archetypical stroke patterns, *lesion atoms*, as resulting from non-negative matrix factorization-based dimensionality reduction. A pattern-discovery framework derived coherent patterns of stroke lesion topographies directly from the segmented high-resolution brain scans from 555 stroke patients. This unsupervised approach led to the derivation of predominantly **right-hemispheric (A) or left-hemispheric (B) lesion patterns.** In case of either one hemisphere, individual *lesion atoms* represented anatomically coherent cortical and subcortical brain regions and respective white matter tracts and had varying emphases on more anterior, medial and posterior regions. In particular, *lesion atoms* 6 and 10 denoted the vascular supply territory of the posterior circulation. While subcortical basal ganglia lesions and cortical lesions in anterior and insular regions were combined in a single *lesion atom* on the left side of the brain, these patterns were characterized by two separate *lesion atoms* on the right side of the brain. **C. Similarity of lesion patterns across patients.** The maximum lesion overlap wa localized subcortically and in proximity of insular regions in the left and right vascular supply territory of the middle cerebral artery. There were neither significant region-wise difference between men and women, nor between left and right hemisphere (**Supplementary Tables 1 and 2**). Some *lesion atoms* did not comprise any substantial contributions from subcortical brain regions and are shown in transparent.

The low-dimensional representation of lesion topographies served as input for fully probabilistic, hierarchical linear regression models to explain acute stroke severity: We here first examined general effects across all patients, on the level of *lesion atoms* and on the level of individual anatomical brain regions. Subsequently, we refined analyses and integrated an additional hierarchy capturing sex-specific effects. We here stratified for male and female status and scrutinized sex-specific effects of *lesion atoms* and anatomical brain regions. Of note, we corrected all of these analyses for the covariates age, sex, stroke lesion volume, white matter lesion volume and relevant comorbidities (atrial fibrillation, hypertension, diabetes mellitus and coronary artery disease). By including an indicator variable of sex in the model, we captured the general non-lesion pattern specific portion of stroke severity difference between men and women.

### Lesion atom and regional relevance for stroke severity

Out of the ten derived *lesion atoms*, six atoms possessed a substantial explanatory relevance for acute stroke severity. This relevance was inferable from posterior distributions of lesion atom parameters that did not substantially overlap with zero. In the right hemisphere, the most relevant *lesion atom* included subcortical regions, i.e. thalamus, putamen and globus pallidum, as well as the brainstem (mean of the posterior distribution=3.8, highest probability density interval of the posterior distribution covering 90%-certainty (HPDI)=1.85-5.66, **Supplementary Figure 1**). In the left hemisphere, the most relevant *lesion atom*, was characterized by both subcortical and cortical regions (posterior mean=3.05, HPDI=2.33-3.77, **Supplementary Figure 1**). Affected left and right subcortical regions were similar, whereas left cortical affected regions additionally included inferior frontal, insular and superior temporal gyrus regions as well as pre- and postcentral gyri.

Once projected back to the level of individual grey matter regions and white matter tracts, similarities and disparities between left and right hemisphere became apparent as well. Subcortical regions, most notably putamen, globus pallidus and several white matter tracts (anterior thalamic radiation, corticospinal tract, inferior fronto-occipital fasciculus, superior longitudinal and uncinate fasciculus) explained higher stroke severity, independent of the lesioned hemisphere (**Figure 2**). Likewise, cortical pre- and postcentral as well as insular and opercular regions explained higher stroke severity in both the left and right hemisphere. In contrast, further cortical effects were more pronounced and more widespread in the left hemisphere. These enhanced left-sided effects mainly related to middle and inferior frontal gyri, as well as superior and middle temporal gyri.

**Figure 2.**
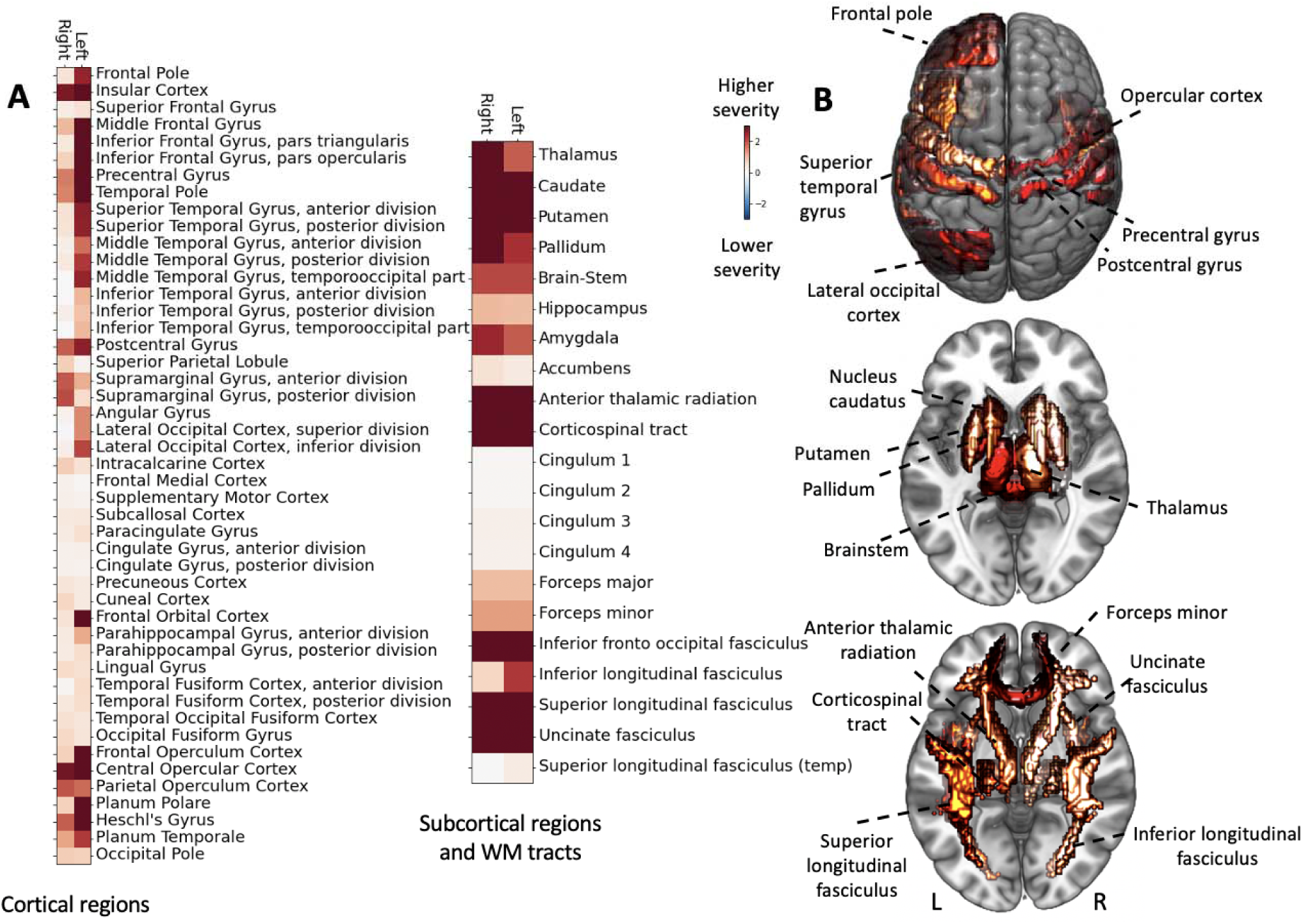
Local brain regions explaining NIHSS-based stroke severity across 555 subjects. **A. Relevant cortical grey matter regions.** Shows collection of marginal posterior parameters from the hierarchical model to explain high vs. low symptom severity (NIHSS). Lesions affecting pre- and postcentral gyri, as well as opercular and insular regions of both hemispheres explained a higher stroke severity. Further brain-behavior effects were left lateralized: Multiple regions including left middle and inferior frontal gyrus as well as superior and middle temporal explained a higher stroke severity only when affecting the left hemisphere. **B. Relevant subcortical grey matter regions and white matter (WM) tracts.** While bilateral subcortical regions in general had substantial effects on stroke severity, the highest weights were assigned to the putamen and caudate, as well as anterior thalamic radiation, corticospinal tract and inferior fronto-occipital fasciculus of the left and right hemisphere.

In summary, we derived stroke severity-linked lesion patterns that highlighted the general importance of subcortical grey matter regions and white matter tracts, as well as of bilateral cortical motor regions, and additional left-lateralized cortical regions, likely underlying language function.

### Differences in lesions patterns between men and women

Subsequently, we concentrated on sex differences in eloquent lesion patterns. We refined our Bayesian model and introduced a hierarchical structure that allowed *lesion atom* effects on stroke severity to vary by sex. Previous findings suggest a higher stroke severity in women in general, the extent of which cannot be sufficiently explained by neither their advanced age, nor differences in lesion volume.^23^

Main patterns of explanatory relevances remained similar to the preceding analysis across all subjects: For both men and women, the same two *lesion atoms*, that had already emerged in the joint analyses, had the highest explanatory relevance. The right-hemispheric *lesion atom* denoted subcortical regions, amongst others representing thalamus, putamen and globus pallidum (men: posterior mean=2.81, HPDI=0.796-4.85, women: posterior mean=3.00, HPDI=0.641-5.3, **Figure 3**, *lesion atom* 3). The lesion atom in the left hemisphere combined the same subcortical structures and additional cortical regions, all of them in proximity to insular cortex in the left hemisphere (men: posterior mean=3.00, HPDI=2.02-3.96, women: posterior mean=2.52, HPDI=1.43-3.49, **Figure 3**, lesion atom 7).

**Figure 3.**
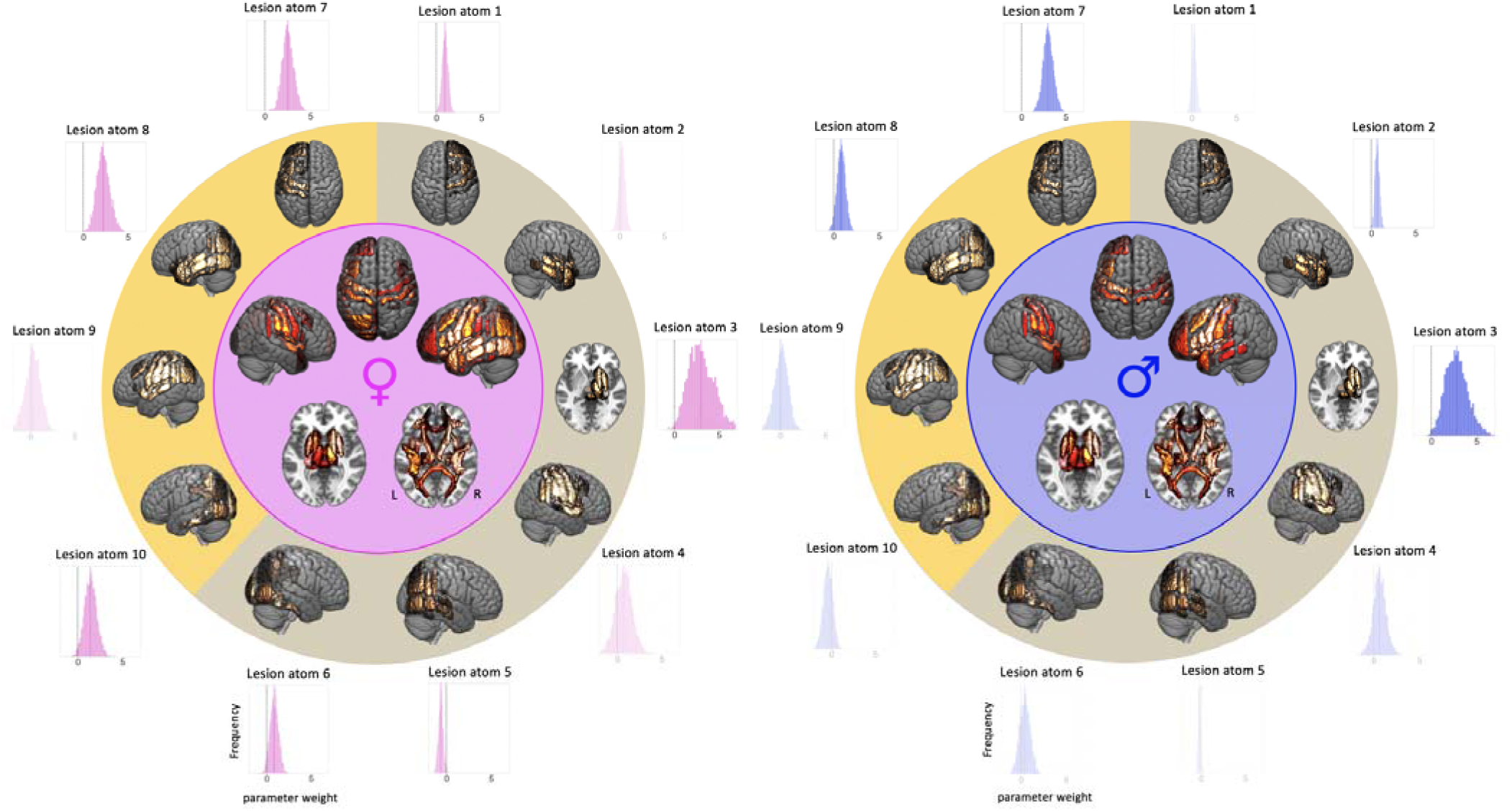
Sex-specific posterior distributions of all ten lesion atom parameters and overall whole-brain region-wise relevance to explain stroke severity in women (*left)* and men (*right*). Our Bayesian framework was purpose-designed to enable fully probabilistic estimation of the parameter that quantify the associations of the ten *lesion atoms* with stroke severity. These posterior parameter distributions are shown in outer circles, corresponding *lesion atom* renderings are presented in the subjacent circle (right-hemispheric *lesion atoms*: shaded in *yellow-olive*, left-hemispheric *lesion atoms*: *orange-yellow*, distributions that substantially diverged from zero are non-transparent). *Lesion atom* 7 of the left hemisphere and *lesion atom* 3 of the right hemisphere had the highest weights, implying a high relevance in explaining stroke severity, in both men and women. In view of seven relevant *lesion atoms* in women (*lesion atoms:* 1, 3, 5, 6, 7, 8, 10), yet only four of such relevant *lesion atoms* in men (*lesion atoms:* 2,3,7,8), lesion patterns were more wide-spread in women compared to men. This female-specific more wide-spread pattern becomes additionally apparent in whole-brain renderings of region-wise relevances explaining stroke severity, as visualized in circle centers (c.f., Figure 4 for details).

**Figure 4.**
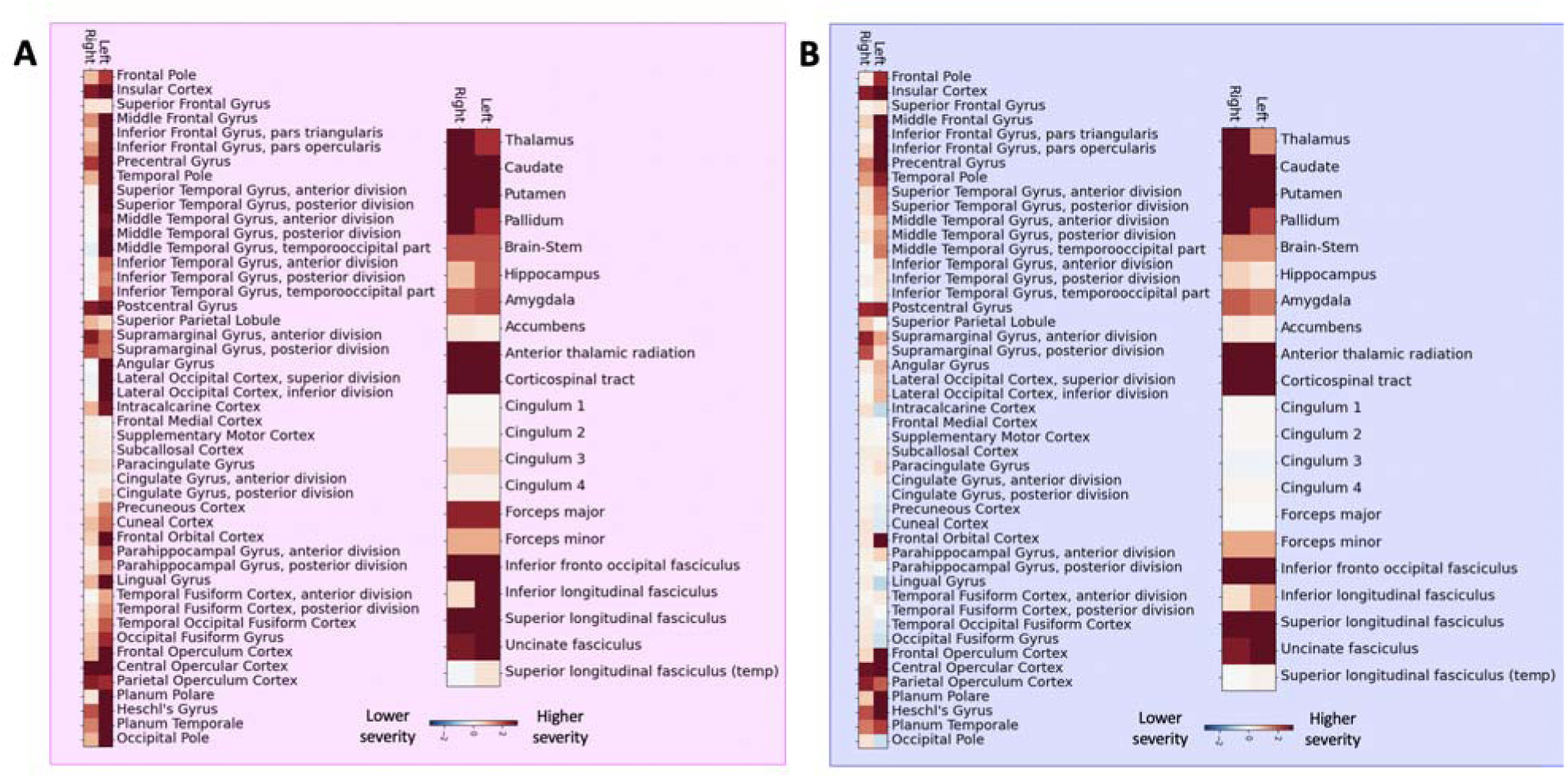
Local brain regions explaining stroke severity in (A) 208 women and (B) 347 men. In both men and women, subcortical lesions affecting grey matter regions and white matter tracts explained higher stroke severity. Similarly, cortical presumptive bilateral motor and left-lateralized language regions (e.g. especially bilateral pre- and post-central gyri, left-sided inferior frontal and superior, middle temporal gyri) also explained higher stroke severity. In difference to men, women featured more widespread and also more pronounced lesion pattern, including a greater range of cortical regions contributing to stroke severity, e.g. the left inferior temporal gyrus, left angular gyrus and lateral occipital cortex, lingual gyrus as well as precuneus and cuneal cortex gyrus. These differences in eloquent lesion patterns arose despite comparable total lesion volumes for men and women (two-sided t-test: *p*>0.05).

Women presented with generally more widespread explanatory relevances as seven out of ten *lesion atoms* posterior distributions did not overlap with zero (**Figure 3**, *lesion atoms* 1, 3,5-8,10). In men, only four *lesion atoms* posterior distributions did not overlap with zero (**Figure 3**, *lesion atoms* 2-3,7-8).

Once projected back to the level of individual brain regions, these more widespread lesion pattern effects in women were also visible, in particular regarding cortical grey matter regions (c.f., **Figure 3** **& 4**). Manifest differences between men and women emerged for two specific *lesion atoms*: For women, *lesion atoms* 1 and 10 had substantial higher explanatory relevances, i.e. the distribution of the difference between posterior distributions of male and female patients did not overlapping with zero (**Figure 5**). *Lesion atom* 1 was mainly characterized by right-sided frontal, insular and opercular as well as pre- and postcentral regions.

**Figure 5.**
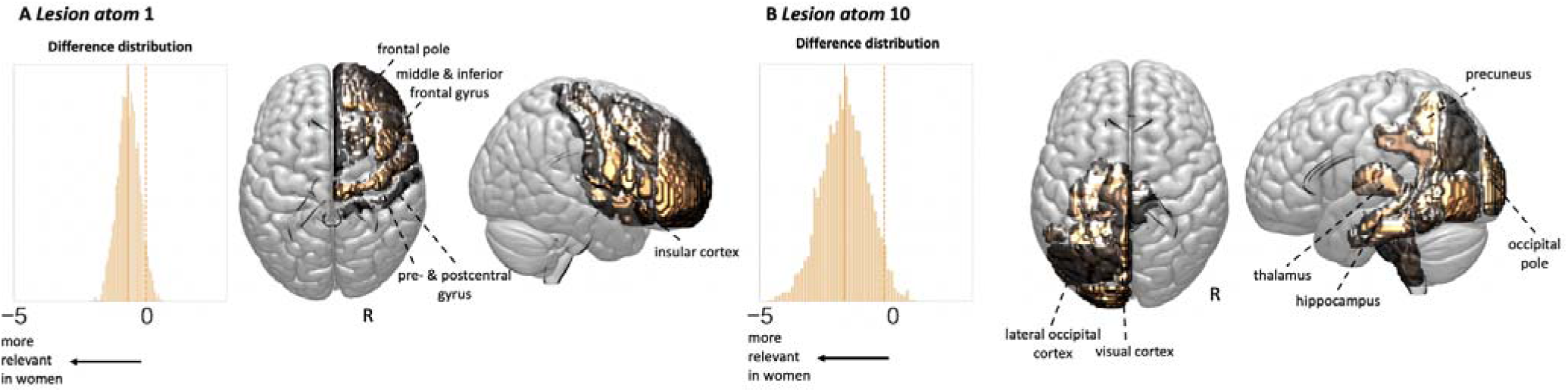
Two lesion atoms showed a substantially stronger stroke-induced symptoms in women than men. **A. *Lesion atom* 1.** Less than 5% of samples of the difference distribution between men and women (i.e., posterior parameter distribution of *lesion atom 1* in men – posterior parameter distribution of *lesion atom 1* in women) overlapped with zero, suggesting a substantially larger lesion atom 1 effect in women. This specific lesion pattern represented right-hemispheric lesions in frontal, insular and pre- and post-central regions (difference distribution: mean=-0.70, HPDI=-1.41- −0.02). **B. *Lesion atom* 10.** Lesions in left-sided brain regions of the posterior circulation caused more severe strokes specifically in women (difference distribution: mean=-1.82, HPDI=−3.16- −0.319). The sex difference for *lesion atom* 10 can be considered more pronounced and robust, given it was reliably observable in ancillary downsampling analyses and was replicated in an independent dataset of stroke patients.

*Lesion atom* 10 primarily represented regions in the left PCA-territory (left hippocampus, thalamus, precuneus & further occipital regions).

### Ancillary analyses

Sex differences in both of these *lesion atoms* 1 and 10 remained observable after downsampling the group of men with milder NIHSS stroke severity, i.e. after harmonizing both the ratio of women-to-men as well as their stroke severity (*p*>0.1 in all cases). *Lesion atom* 1 comprised significant sex differences as indicated by difference distributions not overlapping with zero in three out of 20 random downsampling analyses. When difference distributions overlapped with zero, the posterior parameter distribution of *lesion atom* 1 still substantially differed from zero exclusively for women, yet not for men in all but one case (**Supplementary Figure 2A**). Even more convincing results, suggesting robust sex differences, were obtained for *lesion atom* 10: In nine out of 20 random downsampling analyses, difference distributions did not overlap with zero and the majority of remaining posterior distributions differed from zero for women only, but not for men (in 7 out of 11 cases, **Supplementary Figure 2B**). In further ancillary analyses, we aimed to explore the effects of sexual hormones, such as estrogen, which are known to be markedly affected by menopause.^24^ We stratified the entire group of patients according to their sex and an age cutoff of 52 years, the median age at menopause.^25^ While any of the sex differences for *lesion atoms* 1 and 10 observed in the main analysis disappeared in the analysis of all men and women below the age of 52 years, these sex-specific effects in *lesion atoms* 1 and 10 became even more apparent when restricting the analysis to all men and women above the age of 52 years. However, no conclusive contrast was found between women above and below the age of 52 years (**Supplementary Figure 3**).

### Replication analyses

Similar main findings were noticed when repeating analyses in a completely independent, multisite dataset.^26^ Once again, we extracted ten *lesion atoms* that captured typical stroke patterns in the left and right hemisphere in low dimensions (**Supplementary Figure 4**). *Lesion atoms* of the development and replication dataset were highly correlated, indicating our unsupervised approach facilitated the recovery of similar lesion topography embeddings in both independent datasets (**Supplementary Table 3**).

The most relevant regions explaining stroke severity were also located subcortically in the left and right hemisphere as well as in bilateral pre- and postcentral gyri and left-hemispheric insular and opercular regions (**Supplementary Figure 5A**). Women, once again, presented more widespread eloquent lesion patterns compared to men (**Supplementary Figure 5B&C**). In particular, we found substantial differences between men and women in a *lesion atom* that predominantly comprised left-hemispheric posterior cerebral artery-supplied regions (difference distribution: mean=-2.68, HPDI=-4.92- −0.733 (i.e. no overlap with zero), *lesion atom* 10). In combination with our ancillary downsampling analyses, sex differences relating to the *lesion atom* characterizing left-sided PCA-supplied regions were the most pronounced and robust.

## Discussion

We here combined a novel probabilistic lesion symptom mapping technique with empirical lesion data originating from two large independent cohorts of 1,058 AIS patients to derive and validate sex-specific lesion patterns underlying NIHSS-based acute stroke severity.

Across all patients, lesion patterns highlighted the relevance of bilateral subcortical and white matter regions, as well as bilateral cortical motor regions and left-lateralized cortical insular and opercular regions, likely representing regions underlying language function. This distribution of main weights was rendered particularly plausible, given that the NIHSS scale assigns the majority of its points to motor and language functions. Additionally, results from both of our cohorts, and those results originating from established voxel-wise lesion symptom mapping analyses^20^ were very similar. This congruency increased the confidence in our methodological approach, as well as in the accuracy and reproducibility of results.

When eventually comparing men and women, eloquent lesion patterns were generally more widespread in female patients, implying that more regions contributed to stroke severity in women. These sex discrepancies were particularly pronounced for lesions affecting left-hemispheric regions of the posterior circulation. Lesions in these regions led to a substantially higher stroke severity in women only, indicating sex-specific distributions of brain function and potentially far-reaching implications for future treatment decisions in men and women.

Discussing ascertained sex-specific findings in further detail: First of all, we could not link sex differences in stroke severity to differences in total normalized lesion volume, or in any normalized lesion volume of the atlas-based brain regions and white matter tracts. Rather, similar lesion patterns appeared to lead to more severe strokes in women only, yet not in men. Stroke severity in men was predominantly explained by four specific lesion patterns, denoting bilateral subcortical and left-sided insular and opercular regions. Women presented with similar eloquent lesion patterns, but were characterized by several further relevant lesion patterns. These additional lesion patterns did not possess any relevance in men, therefore, the female-specific effects can be considered more spatially widespread. Most noticeably, lesions in the vascular territory of the *left* PCA, amongst other regions affecting left hippocampal, thalamic regions, left fusiform, lingual and intracalcarine cortex as well as left precuneus and cuneal cortex, explained a disproportionally high stroke severity in women than men. This difference was stably observable across datasets and did not appear to represent an artifact of slightly greater stroke severity in women or imbalanced numbers of men and women in our sample.

It is important to note that the strongest sex differences, that we observed, were strictly lateralized to the *left* hemisphere. Previous research suggests that male or female sex and respective sex hormones contribute to induce functional cerebral asymmetries.^27^ Men appear to have a stronger hemispheric asymmetry; however, while robustly replicated, determined effect sizes have been very small.^28^ Such an enhanced asymmetry in men was also found in some early lesion studies on intelligence.^29^ However, further early lesion studies suggest that lateralization differences between the sexes might be even more complex: Inglis and colleagues underline that female brains may be asymmetric to a comparable degree, yet in different ways.^30, 31^ In particular, they found that left-hemispheric lesions in women led to both verbal and performance scale IQ deterioration, while only one quality – either verbal or performance – was affected in all other lesion and sex constellations.^30, 32^ Our outcome measure, NIHSS-based stroke severity, cannot be broken down and thus does not allow for conclusions on very specific functions, such as verbal and performance scale IQ. This level of granularity hampers a direct comparison to earlier studies. Nonetheless, we also find that particularly women are vulnerable to *left-hemispheric* lesions. Indeed, we can relate this excess vulnerability of female versus male patients to anatomically precise lesion locations in left-hemispheric PCA-territory, e.g., featuring hippocampal, thalamic and precuneal regions. Deduced from the existing knowledge on these regions’ physiological functions, we may suppose lesions in these regions more likely underlie (higher) cognitive, than, for example, basic motor functions. This predilection of specific anatomical regions within the left hemisphere could also explain differences that arose in earlier lesion studies as these studies stratified patients only based on left- or right-sided lesion locations without appreciating any greater spatial detail of lesions.^29, 30, 32^

Altogether, a sex-specific lateralization effect, comparable to the considerable extent and spatial distribution of the one that we detected in two independent datasets, has neither been described in previous lesion studies, nor studies focused on sex discrepancies in functional cerebral asymmetries in healthy adults.^28^ Conceivably, this discrepancy between previous reports and our findings may originate from i) employing the just mentioned greater anatomical resolution in our study compared to early sex-aware lesion studies, ii) not explicitly considering male or female phenotype in any of the recent lesion symptom studies^17, 20, 33, 34, 35^ and iii) the fact that our lesion symptom analysis jointly investigated effects of functional asymmetry and the capacity for acute compensation, e.g., by means of brain plasticity.

Intriguingly, we furthermore witnessed signs of an interaction effect of sex with age, when stratifying the entire sample based on the median age at menopause.^25^ While there was no difference between men and women below the age of 52 years, the difference relating to the left-hemispheric PCA-territory was even more pronounced in men and women above the age of 52 years. This constellation was thus suggestive of a critical influence of sexual hormones, that are dramatically changed in women after menopause.^36^

Altogether, sex differences in brain organization in general and in stroke incidence and outcome in particular are to a large extent attributed to the influence of sex steroid hormones, with estrogen likely being the most prominent one.^36, 37^ These hormones are assumed to unfold their effects via irreversible organizational and reversible activational mechanisms. The former effect implies the facilitation of definite male or female tissue phenotypes, while the latter requires the presence of the hormone for an effect.^24^ Any sex difference that changes with menopause is thus rather caused via activational hormonal pathways. An example of sex differences in stroke thought to be due to activational hormonal effects is the reduced stroke risk due to the pre-menopausal cycle of estrogens. This protective effect is lost after menopause^38^ and could – so far – not be reestablished by hormone replacement therapy independent of the time of initiation.^39, 40^ Experimental research has furthermore shown that female animals experience smaller stroke lesions than male animals.^41^ This effect could be neutralized by ovariectomy and consequently a decrease in estrogen levels.^42^ It was interpreted as hormone-linked sex-specific sensitivity to cerebral ischemia.^24, 43^ In fact, male-specific cell cultures of hippocampal neurons and astrocytes seem to be more vulnerable to ischemia than female-specific cell cultures^44^ and even ischemia-independent research indicates an important role of estrogen in sustained hippocampal structural plasticity and associated cognitive function.^45, 46^

The effect of age that we witnessed here – stronger sex-specific effects in older patients and no sex-specific effects in younger patients – also hints at an activational nature of the apparent sex disparities. While our outcome measure of global stroke severity does not allow for fine-grained evaluations of implicated functions, let alone hippocampus-linked cognitive functions, it is nonetheless notable that hippocampal regions, that were markedly susceptible to hormonal influences as outlined above, contributed to the female-specific lesion patterns uncovered in our study.

Independent of their exact origin, the sex-sensitive effects, that we observed in brain-behavior associations underlying stroke severity, entail important clinical implications. Since lesions of any kind explained more severe stroke in women, rescuing the same (normalized) amount of brain tissue – for example by thrombolysis or mechanical thrombectomy – is likely to have a more beneficial effect in female than male patients. This resulting expectation is well in line with previous reports on enhanced therapy response to intravenous thrombolysis^47, 48, 49^ or more advantageous long-term outcome in women compared to men in clinical mechanical thrombectomy studies.^50^ Interestingly, the just recently published study on sex discrepancies after thrombectomy by Sheth and colleagues ascertains these outcome differences between men and women despite comparable infarct volumes and reperfusion rates.

Most previous studies have, however, relied on sex-independent, general cutoffs of salvageable tissue volumes to, for example, decide upon whether to undertake mechanical thrombectomy in the later time window.^51^ Future clinical treatment studies could therefore systematically test varying cutoffs for men and women, hypothesizing that rescuing a lesser amount of tissue in women would still be sufficient for a noticeable positive treatment response.

Furthermore, it may be particularly essential to take into account this female-specific salvaging effect for lesions in the posterior circulation territory of the left hemisphere and future thrombectomy studies could evaluate, whether female patients benefitted more from endovascular reperfusion of more distal PCA-occlusions.

Consequently, a very effective step towards tailored medical stroke care^52, 53^ may simply lie in more sex-aware acute treatment decisions. Sex-specific guidelines on stroke prevention^7^ could be complemented by sex-specific guidelines on acute treatment decisions; enhancing sex-sensitive stroke care and ultimately increasing the benefit for both men and women.

### Limitations and future directions

We did not have any access to information on the hormonal status in women. Instead, we here assumed the same median age of menopause for all female patients. Employing the actual age at menopause, as well as increasing the number of younger patients in general would have allowed for purer conclusions on the potential organizational or activational character of findings. After all, the availability of the exact level of estrogens would have been most desirable, as previous studies indicate that sex-specific functional cerebral asymmetries may even vary during the menstrual cycle.^27^ Therefore, future studies could attempt to gather data on hormonal status and also to recruit more balanced numbers of men and women of pre- and postmenopausal status. Additionally, it might be promising to test links between implicated brain regions and sex-specific genetic underpinnings, as it was recently introduced for healthy population samples^54, 55^

Furthermore, we here explored sex disparities in lesion patterns of *global* stroke severity, which already allows for conclusions on broad clinical implications of our findings. Nonetheless, future studies could immerse into sub-items of the NIHSS or behavioral tests evaluating more specific brain functions at acute and chronic stages. This would be a promising approach to trace back our most prominent sex-specific finding, the relevance of the left PCA-territory, to specific brain functions and explain it in further depth. Incorporating outcomes from more chronic time points would furthermore allow for more definite conclusions on more long-term effects and for example their socioeconomic meaning.

Similarly, the spatial resolution of our Bayesian hierarchical approach stays within the realms of typical lesion patterns and backtransformed atlas brain regions. While a focus on individual atlas regions and potentially even subregions within these atlas regions may be most attractive, it is important not to neglect spatial biases in any spatially more precise, likely voxel-wise, univariate analysis.^56^

Lastly, while we did account for inter-individual differences in important comorbidities, such as atrial fibrillation, and the total volume of white matter hyperintensity lesions, we did not investigate any interactions of these variables with the patient’s sex status. Since first evidence suggests sex-specific effects of white matter integrity on stroke outcome,^57^ future studies are warranted to include not only acute, yet also chronic markers of brain health in a sex-aware manner.

## Conclusions

Stroke severity is often reported to be more severe in women than men. Previous methodology did not allow the evaluation of whether lesion patterns led to these sex differences in outcomes. By deriving a low-dimensional lesion representation and employing Bayesian hierarchical modelling, we here uncovered considerable sex discrepancies in lesion patterns explaining stroke severity. While bilateral subcortical and left-hemispheric inferior frontal, superior and middle temporal regions, i.e., presumed motor and language regions, explained more severe strokes in both men and women, effects in women were more widespread and similar lesions underlay more severe strokes in women compared to men. In particular, lesions in the posterior circulation of the left hemisphere led to a higher stroke severity exclusively in women. These differences were robustly replicated in a second independent, international multi-site lesion dataset and could not be explained by sex differences in lesion volume or more severe stroke in women in general. Altogether, these findings may enhance acute stroke care by motivating more sex-sensitive treatment decisions, for example female and male-specific cut-offs of salvageable tissue for the administration of thrombolysis and thrombectomy. Thereby, our findings hold promise to represent a crucial step towards individualized medicine.

## Methods

### Participant recruitment

Acute ischemic stroke (AIS) patients (n=555), considered as development cohort in this study, were admitted to Massachusetts General Hospital and enrolled as of part the Genes Associated with Stroke Risk and Outcomes Study (GASROS; mean age (standard deviation (SD)): 65.0 (14.8) years, 37% females, c.f., **Table 1** for sex-specific numbers).^58, 59^ Inclusion was generally considered for any AIS patient that met the following criteria; i) adult patients ≥ years of age, ii) admitted to the emergency department with signs and symptoms of AIS, and iii) neuroimaging confirmation of an acute infarct. Only subjects with MRI data obtained within 48 hours from stroke onset as well as complete data on stroke severity and stroke risk factors were included in this study (i.e., complete case analyses, c.f. **supplementary materials** for a sample size justification). Subject gave written informed consent in accordance with the Declaration of Helsinki. The study protocol was approved by the local Institutional Review Board.

### Stroke patient characteristics and imaging

Patients were examined by trained, board-certified vascular neurologists. The recorded sociodemographic and clinical variables included age, sex, and common vascular risk factors (hypertension, diabetes mellitus type 2, atrial fibrillation (AF), coronary artery disease (CAD)). Stroke severity was captured by the acute NIHSS (0: no symptoms, 42: maximum stroke severity).

Each patient was scanned within 48h of admission, standardized clinical imaging protocols included DWI (in the majority of cases on 1.5T General Electric Signa scanners, and a few cases on 1.5 or 3T Siemens scanners, repetition time (TR) 5,000 ms, minimum echo time (TE) of 62 to 117 ms, field-of-view (FOV) 220 mm field-of-view, 5-mm slice thickness with a 1-mm gap, and 0 s/mm^2^ (b-zero) and 1000 s/mm^2^ b-values) and axial T2 FLAIR images (TR 5,000 ms, minimum TE of 62 to 116 ms, TI 2,200 ms, FOV 220–240 mm). Ischemic tissue lesions were manually outlined using semi-automated algorithms.^60^ Raters were blinded to clinical outcomes.

### Magnetic resonance imaging: Preprocessing

Individual images were spatially normalized to standard Montreal Neurological Institute (MNI-152) space: We first linearly realigned both DWI and FLAIR images with an MNI template. Subsequently, we co-registered the DWI image to the FLAIR image, denoised the images^61^ and lastly employed the unified segmentation algorithm to non-linearly normalize the FLAIR image after masking lesioned tissue.^62^ The same transformation was applied to the DWI image as well as the corresponding binary lesion mask. The quality of normalized lesion masks was carefully inspected by two experienced raters (A.K.B, M.B). Insufficient quality, predominantly arising from moderate to severe motion artifacts and/or moderate to severe normalization errors led to the exclusion of subjects (c.f. **supplementary materials** for details). The volume of white matter hyperintensity lesions was obtained via a previously developed, fully automated deep learning-based segmentation pipeline of the white matter hyperintensities on FLAIR images.^63^

Following previous work,^21^ we successively combined 1) the automated low-dimensional embedding of high-dimensional lesion information, and 2) probabilistic modeling to explain the acute stroke severity, as measured by the NIHSS. While we initially determined eloquent lesion patterns across all subjects, we subsequently refined analyses to investigate sex differences on different levels of our Bayesian hierarchical model.

### Derivation of a low-dimensional lesion representation

In view of the high-dimensional lesioned voxel space, we first captured the number of lesioned voxels within 109 brain regions as defined by the Harvard-Oxford atlas.^64^ More precisely, these brain regions represented 47 cortical and 7 subcortical grey matter brain regions per hemisphere, as well as the brainstem. White matter damage was read out in form of lesion load per 20 John-Hopkins-University (JHU)-atlas defined white matter tracts (7 hemisphere-specific tracts, 6 bilateral tracts).^65^ Thus, these steps resulted in 129 parcels in total. To eventually extract reoccurring, archetypical lesion patterns in our stroke population, we performed NMF^22^ on the log-transformed brain region- and tract-wise lesion load. By these means, we obtained ten *lesion atoms*. Advantages in employing NMF may be seen in i) the enhanced interpretability of estimated features, ii) NMF’s multivariate nature, which, importantly, may decrease the distortion of functional localization compared to voxel-wise approaches.^19, 56^

### Explaining inter-individual differences in acute stroke outcomes

These ten NMF-derived lesion atoms, containing information on individual lesion patterns, served as neuroimaging-derived input to our Bayesian hierarchical linear regression model ^66^ to explain acute stroke severity. As in previous work, we aimed to obtain fully probabilistic model parameter estimates that could inform us about each lesion atom’s influence on the outcome. While we first computed the impact of lesion atoms across all subjects in a first Bayesian model, we refined analyses in a subsequent step and introduced a hierarchical lesion atom structure for a second model. This new hierarchy allowed the stratification for an individual’s sex, i.e., we could estimate the lesion atom influence on the outcome in women and men separately. Hence, we obtained separate lesion patterns for men and women.

Prior to carrying out the Bayesian model, lesion atom data, as well as the stroke severity outcome score were corrected for lesion volume.^67^ In addition, our model took into account the effects of (normalized) age, age^2^, sex, and the presence of the following comorbidities: hypertension, diabetes mellitus type 2, atrial fibrillation, coronary artery disease and lastly the log-transformed white matter hyperintensity lesion volume. Importantly, we included sex as a variable in the model to capture sex differences in stroke severity that were *independent* of lesion patterns. If, for example, stroke severity was generally higher in women, without any link to the actual lesion distribution, this effect would be represented by the posterior parameter of this sex variable, yet not in the sex-specific lesion atom posterior parameter estimations. In contrast, sex-specific lesion atom parameters would indicate interaction effects on the outcome, i.e. effects that were specific to an individual’s sex *and* the precise lesion atom.

The full Bayesian hierarchical model specification was as follows:

#### Hyperpriors

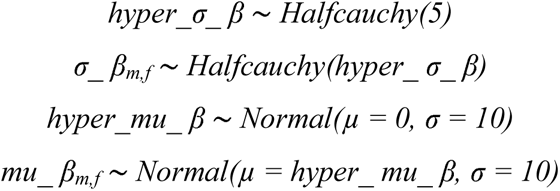

#### Priors

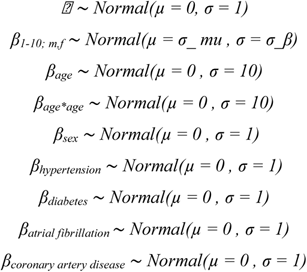

#### Likelihood of linear model

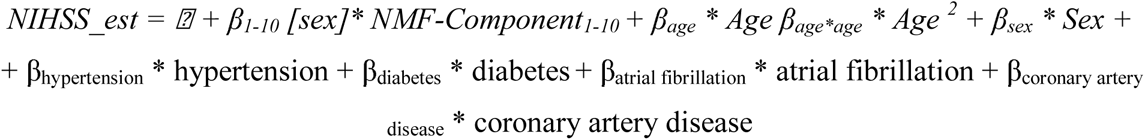

#### Model likelihood

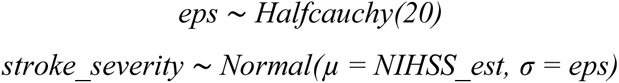

We employed the No U-Turn Sampler (NUTS), a kind of Monte Carlo Markov Chain algorithm (setting: draws=5000),^68^ to draw samples from the posterior parameter distributions.

### Ancillary analyses

Our patient sample comprised a mild male-female imbalance (63% male, 37% female patients) and a higher mean NIHSS score for females (NIHSS_men_: 4.7, NIHSS_women_: 5.6). Therefore, we tested the robustness of our findings by repeatedly downsampling the larger group of men to the same size and comparable stroke severity as the group of women (c.f. **supplementary materials for details**). Additionally, we aimed to explore whether possible sex differences were more likely to be of organizational or activational hormonal nature. Since we did not have access to the precise hormonal status in women, we stratified our sample based on an age cutoff of 52 years, according to the median age at menopause.^25^ Therefore, we repeated main analyses in i) a sample of women only (below versus above 52 years of age, 41 younger and 167 older patients), ii) men and women below the age of 52 years (71 male and 41 female patients), iii) men and women above the age of 52 years (276 male and 167 female patients).

### Replication analysis

We repeated analyses in a completely independent dataset of 503 ischemic stroke patients (age: 65.0 (14.6), sex: 40.6% female, NIHSS: 5.48 (5.35)), acquired within the framework of the multi-site MRI-GENIE study,^26^ to test the robustness of our findings (c.f. **supplementary materials for details**).

### Code availability

Preprocessing of MRI images was conducted in a Matlab 2019b framework (The Mathworks, Natick, MA, USA), the packages Statistical Parametric Mapping (SPM12; http://www.fil.ion.ucl.ac.uk/spm/) and the ancillary package SPM_Superres (https://github.com/brudfors/spm_superres). Further analyses were implemented in Python 3.7 (primarily packages: nilearn^69^ and pymc3^70^). Full code for reproducibility and reuse is available here: www.github.com/TO_BE_ADDED

## Abbreviations

AIS: acute ischemic stroke
MCA: middle cerebral artery
NIHSS: National Institutes of Health Stroke Scale
PCA: posterior cerebral artery

## Acknowledgements

We would like to acknowledge valuable comments by Parashkev Nachev, M.D., Ph.D. on previous versions on this manuscript.

## Supplementary materials

### Development: GASROS

#### Sample size derivation

We had access to 668 patients with manual lesion segmentations. 579 out of these 668 patients had complete data with respect to clinical variables (i.e., age, sex, stroke severity, comorbidities, WMH lesion volume). Quality control of normalization results of structural images led to the secondary exclusion of 24 out of these 579 patients, leaving 555 patients for final analyses.

**Supplementary Figure 1.**
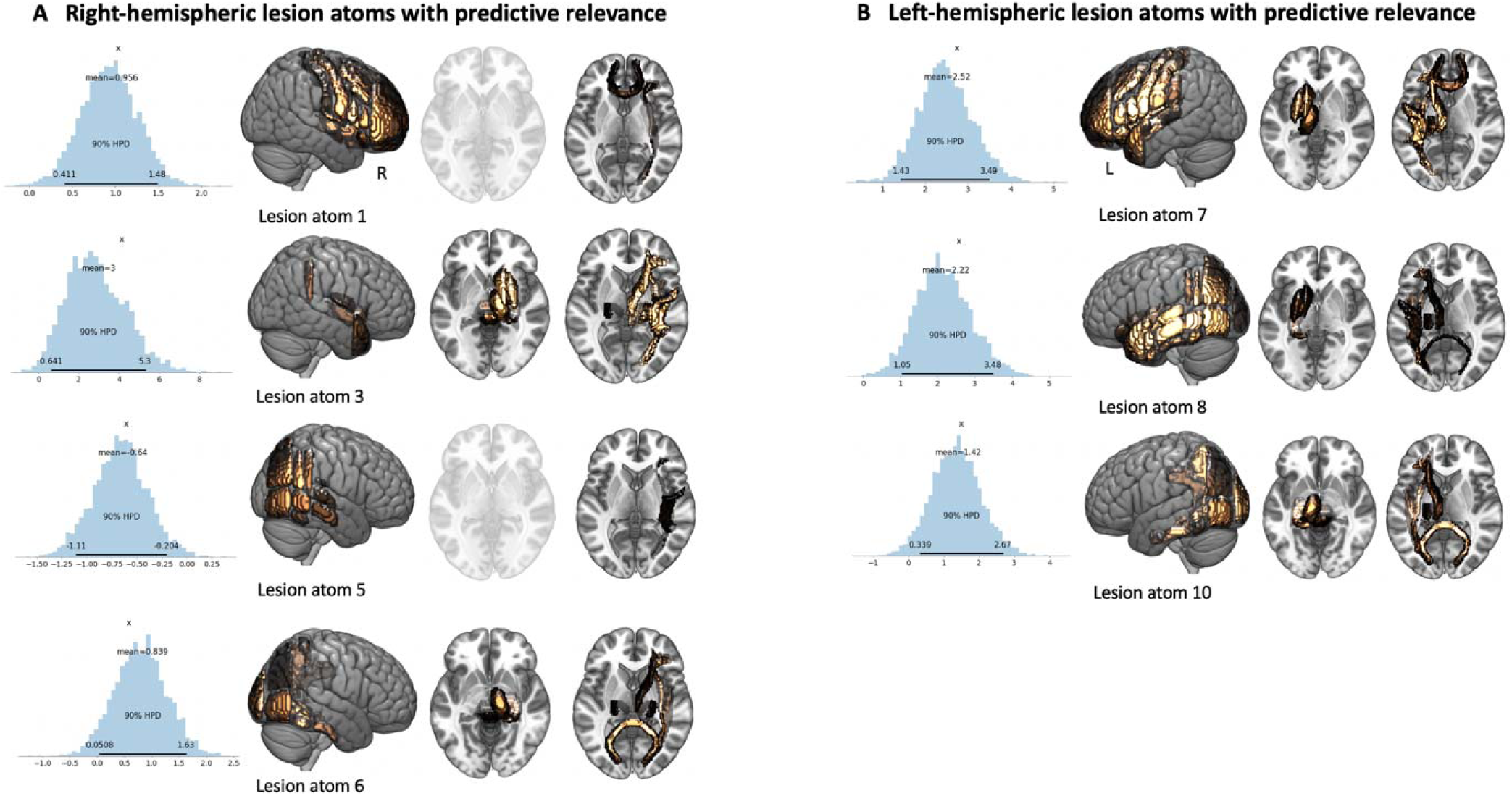
Posterior distributions of lesion atom parameters that substantially diverged from zero. Right-hemispheric *lesion atom* 3 and left-hemispheric *lesion atom* 7 presented the highest predictive relevances of stroke severity.

#### Ancillary analyses: Downsampling

To harmonize the entire sample with respect to the number of included men (n=347, 63%) and women (n=208, 37%) and their acute stroke severity (mean NIHSS_men_: 4.7, NIHSS_women_: 5.6, *p*=0.09), we downsampled the larger group of men. We did so proportional to the number of women with i) NIHSS scores between 0 and 9 (n=167) and ii) those with NIHSS >9 (n=41). That is, we chose we randomly chose 167 out of 289 male patients with an NIHSS between 0 and 9 and 41 out of 58 for an NIHSS>9. We repeated this downsampling step 20 times. In the following, we performed the same Bayesian hierarchical modelling analysis as outlined for the entire sample (Methods: Predicting inter-individual differences in acute stroke outcomes). We specifically evaluated the stability for the main findings of sex-specific relevances of *lesion atoms* 1 and 10 in these 20 analyses.

**Supplementary Figure 2A.**
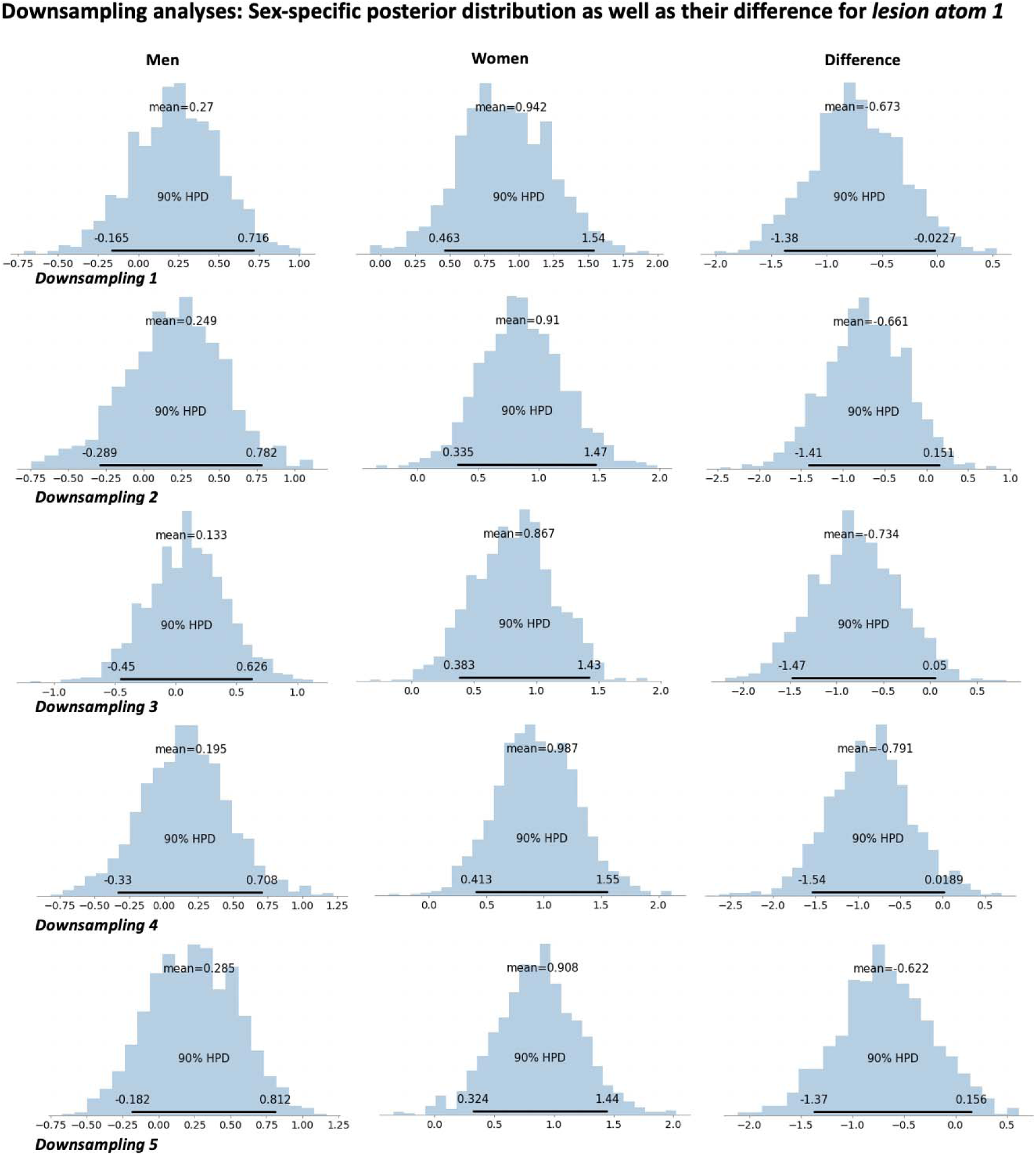
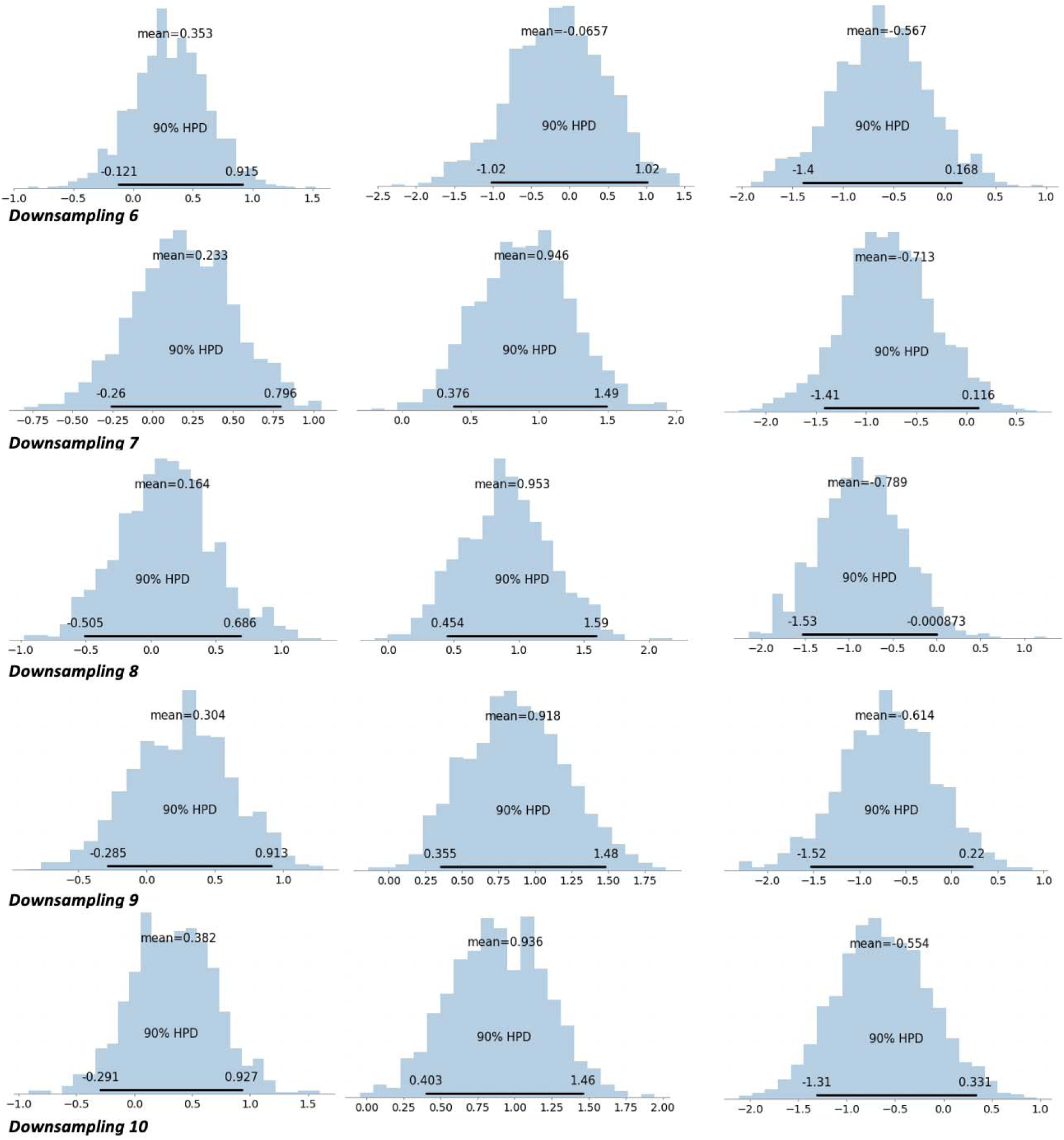
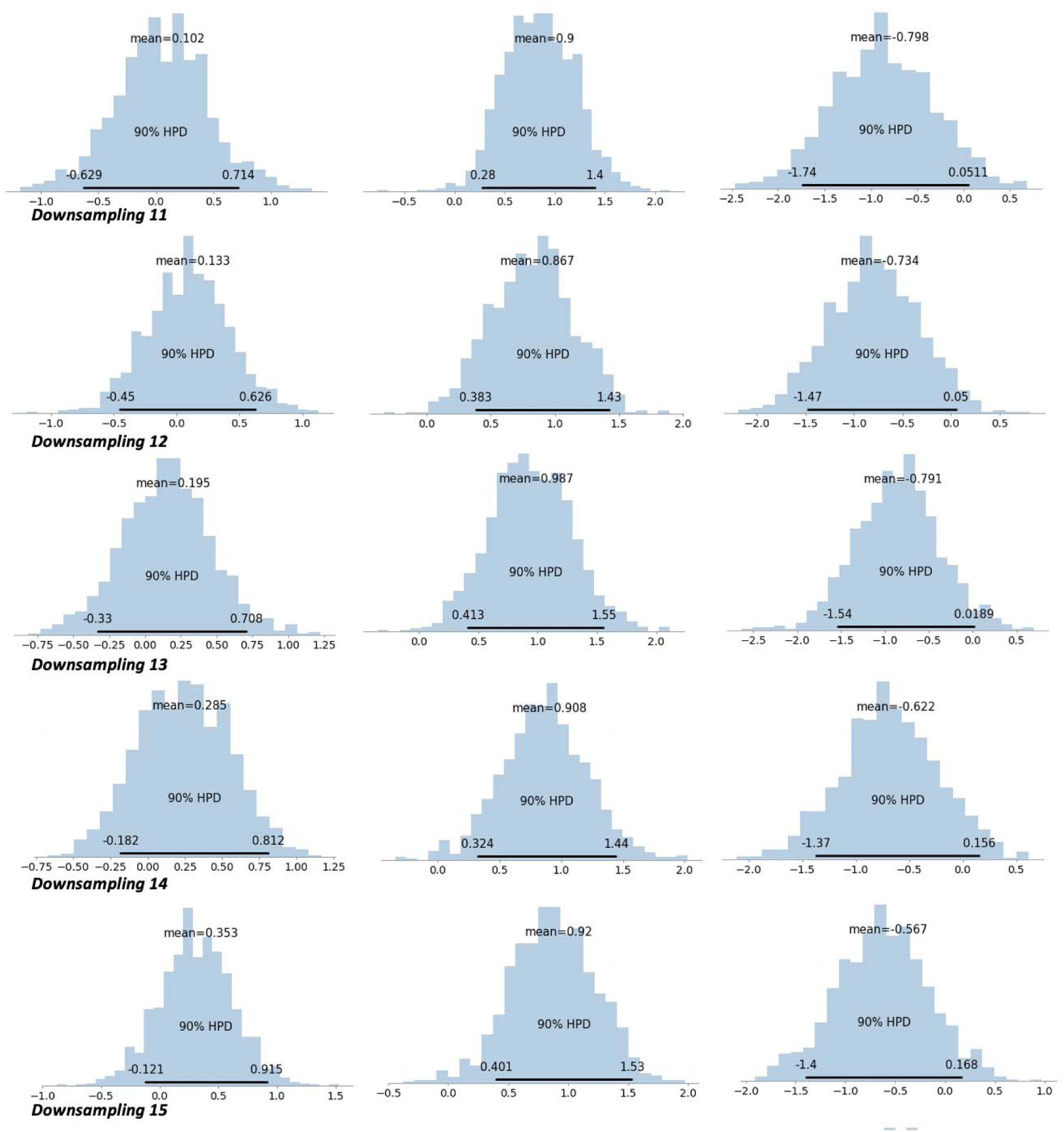
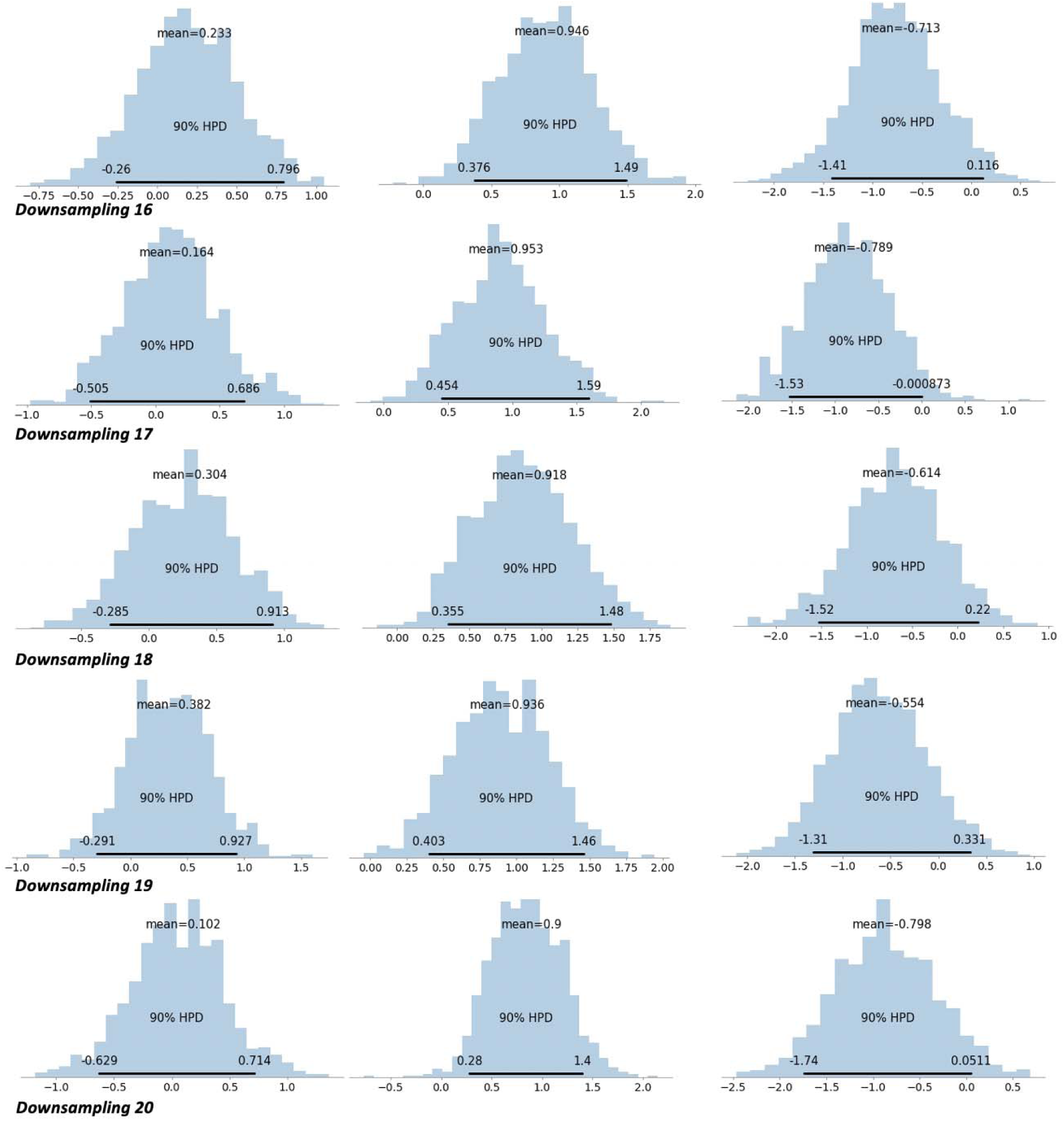
Downsampling results for *lesion atom* 1.

**Supplementary Figure 2B.**
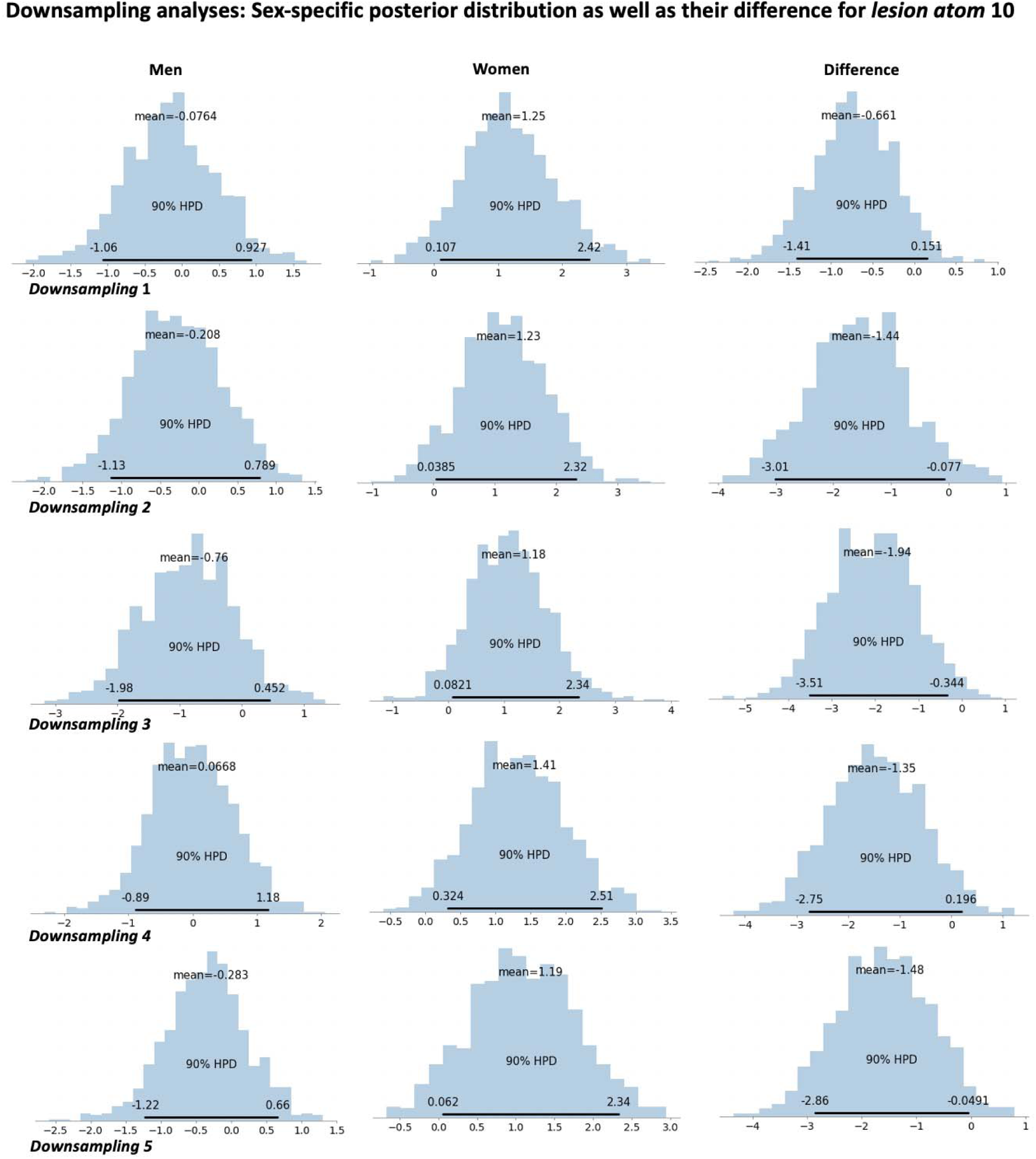
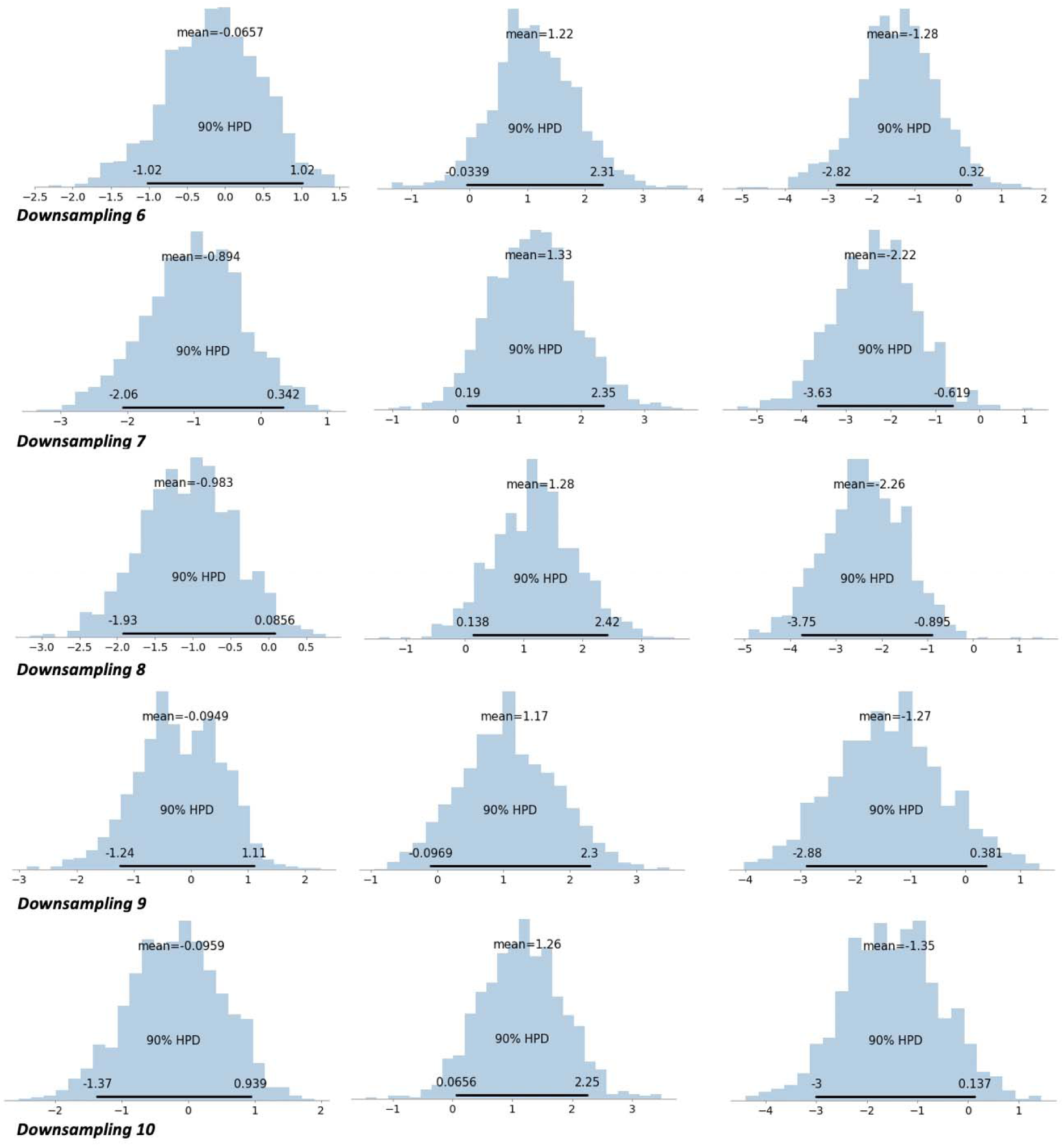
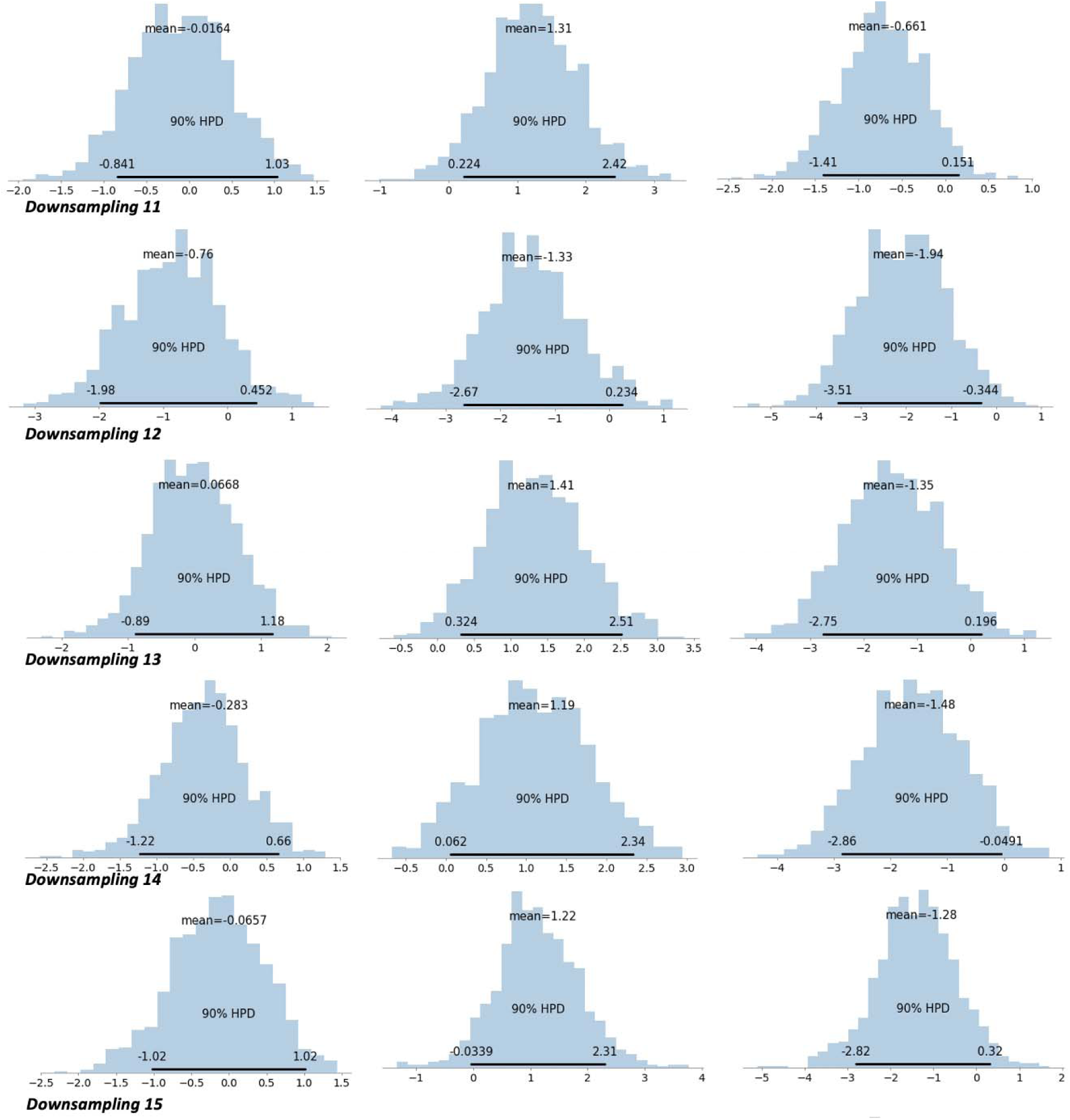
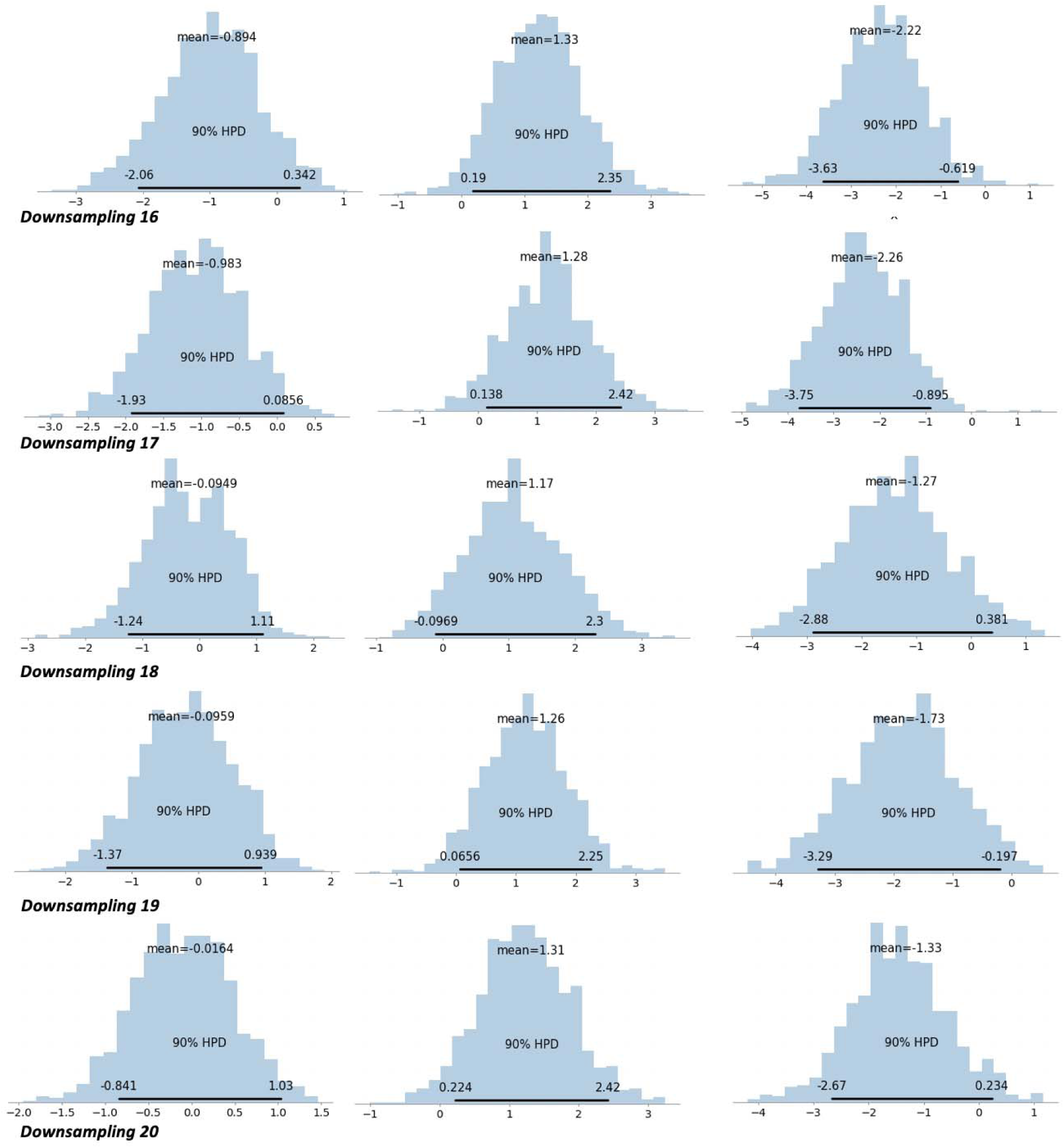
Downsampling results for *lesion atom* 10.

**Supplementary Figure 3.**
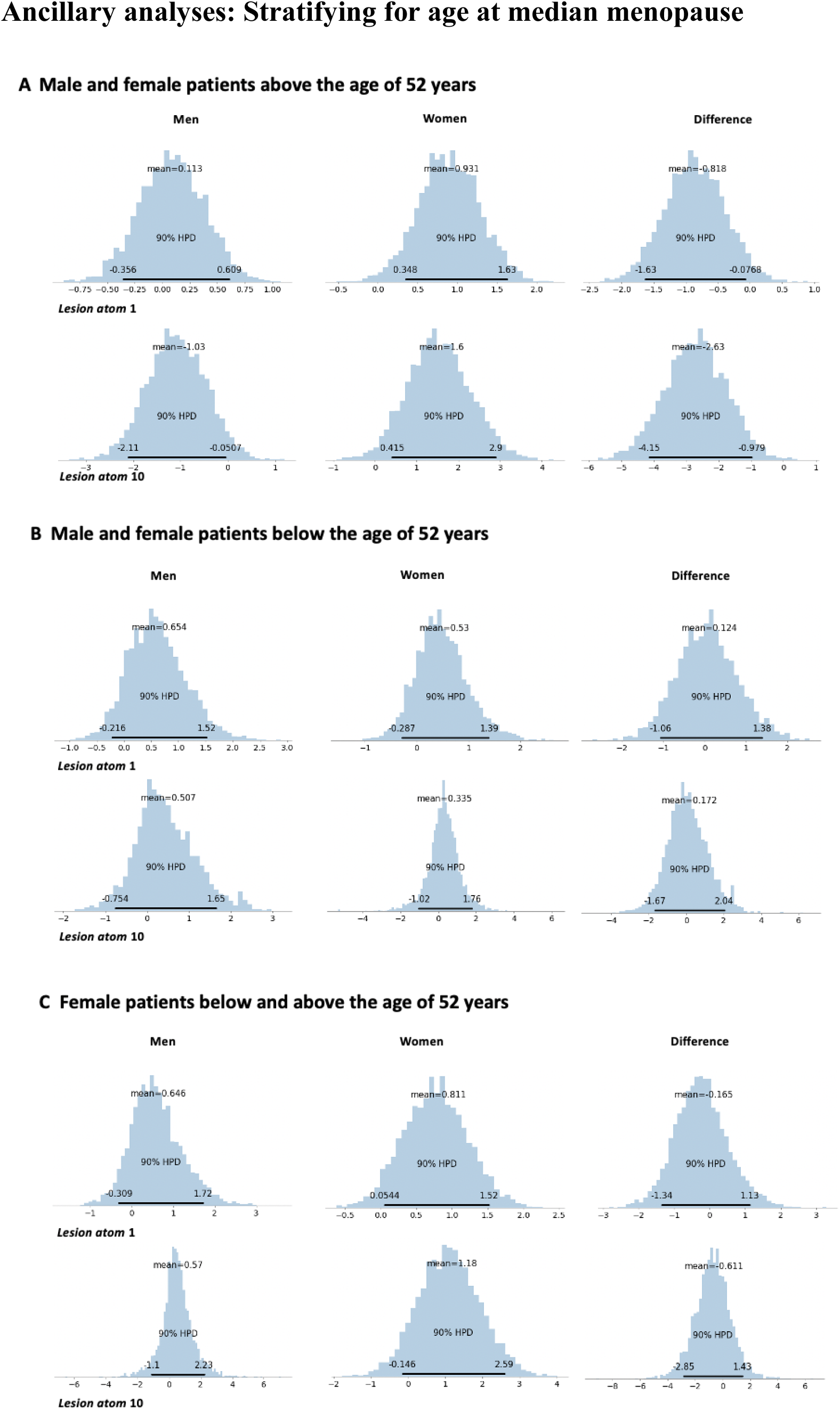
Posterior parameter distributions for *lesion atoms* 1 and 10 that featured substantial sex-differences in the main analysis. These differences were pronounced in the sample of older subjects (above median age of menopause, A), yet not visible in the sample of younger subjects (B) and women only (C).

### Replication: MRI-GENIE

AIS patients in the replication dataset originated the MRI-GENIE study (Giese *et al.*, 2017). Out of 2,765 automatically segmented lesions (Wu *et al.*, 2019), 1,920 (70.1%) passed internal quality control by two raters (M.B., A.K.B.). Included and excluded patients did not differ with respect to age, sex, NIHSS stroke severity and Rankin Scale-based functional outcome (*p*>0.05, Bonferroni-corrected for four comparisons). Lesions were spatially normalized to MNI-space (Wu, 2019).

Initial NIHSS-based stroke severity was available for 942 MRI-Genie patients from six international centers. We excluded those subjects that were enrolled in the GASROS study to prevent an overlap of data between the development and replication cohort (n=150). Automatically segmented lesion outlines were available for 503 out of the remaining 792 patients with information on stroke severity. Thus, these 503 patients (age: 65.0 (14.6), sex: 40.6% female, NIHSS: 5.48 (5.35)) originating from five centers constituted the finally included sample. Subject gave written informed consent in accordance with the Declaration of Helsinki. The study protocol was approved by the local Institutional Review Board.

### Low-dimensional lesion embedding via non-negative matrix factorization

We once again estimated ten lesion atoms that represented typical voxel-wise lesion pattern. The derived lesion atoms could be matched with those atoms estimated for the data in the development cohort, which facilitated the comparison of results further (**Supplementary Table 3**).

**Supplementary Figure 4.**
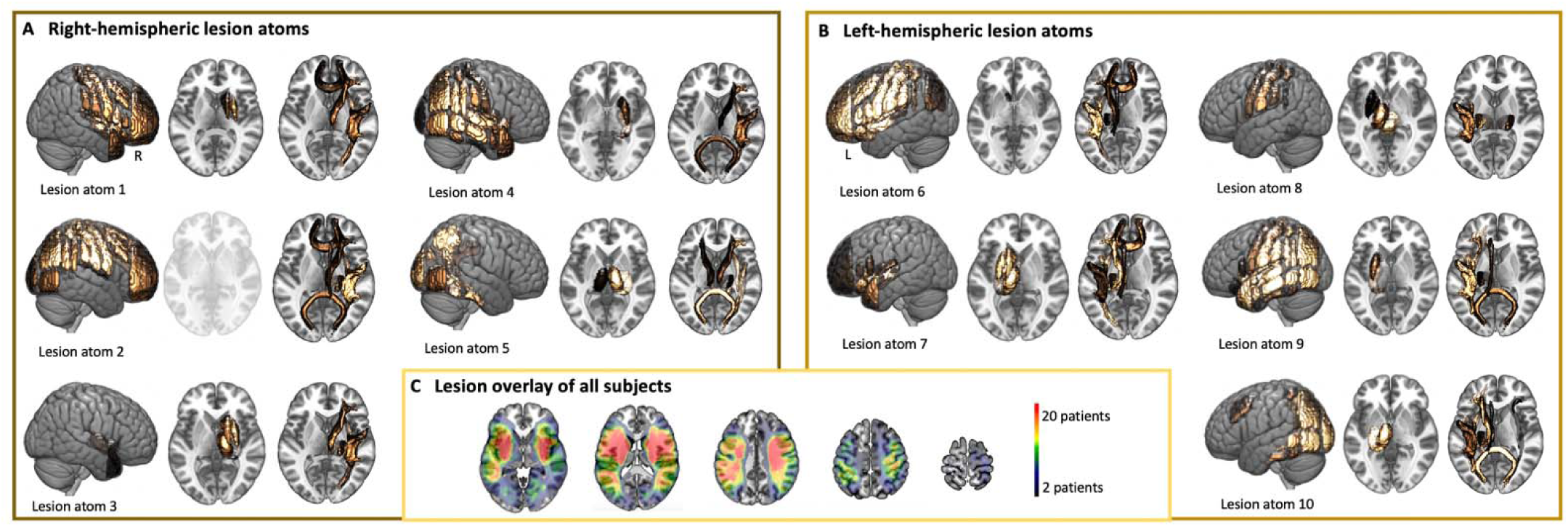
Low-dimensional lesion representation in MRIGenie.

**Supplementary Figure 5.**
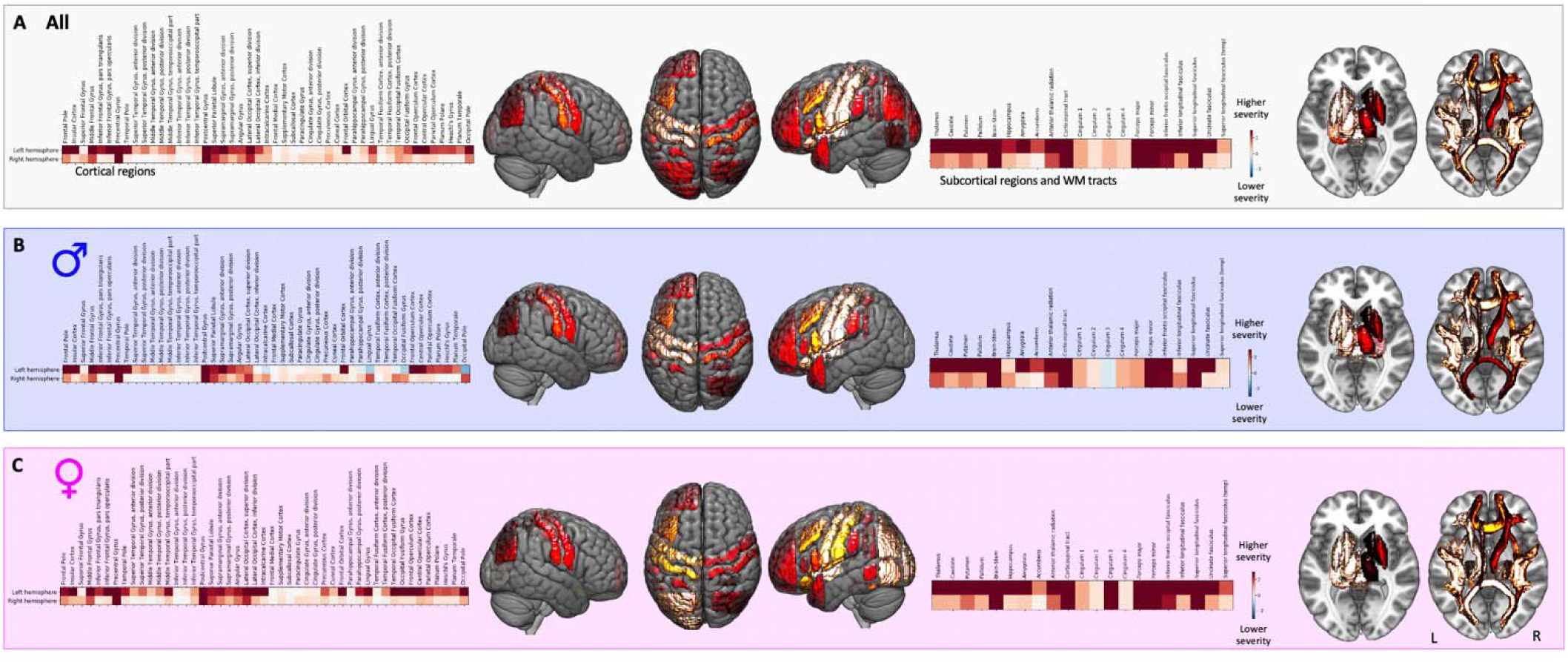
Predictive relevances of individual brain regions for all (A), male (B) and female patients (C). Once again, female patients featured a more wide-spread pattern, particularly comprising brain areas in the posterior circulation.

**Supplementary Table 1.**
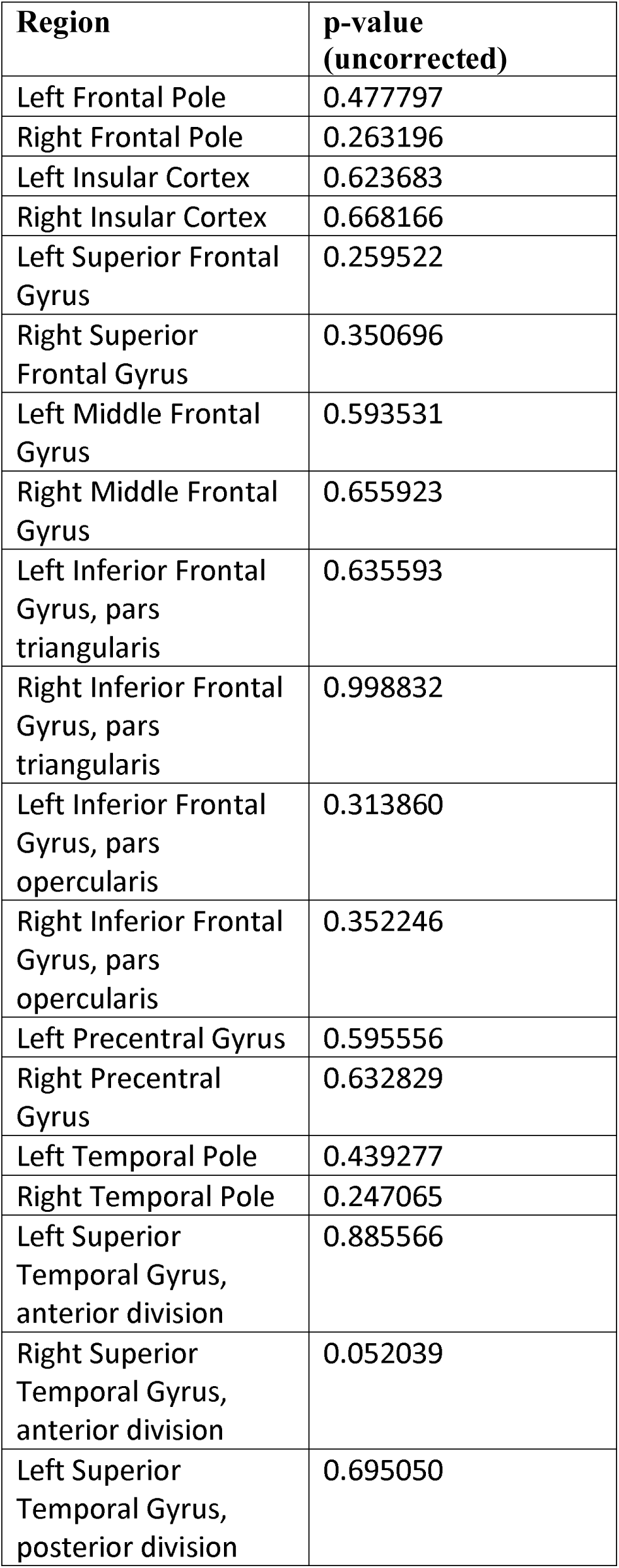
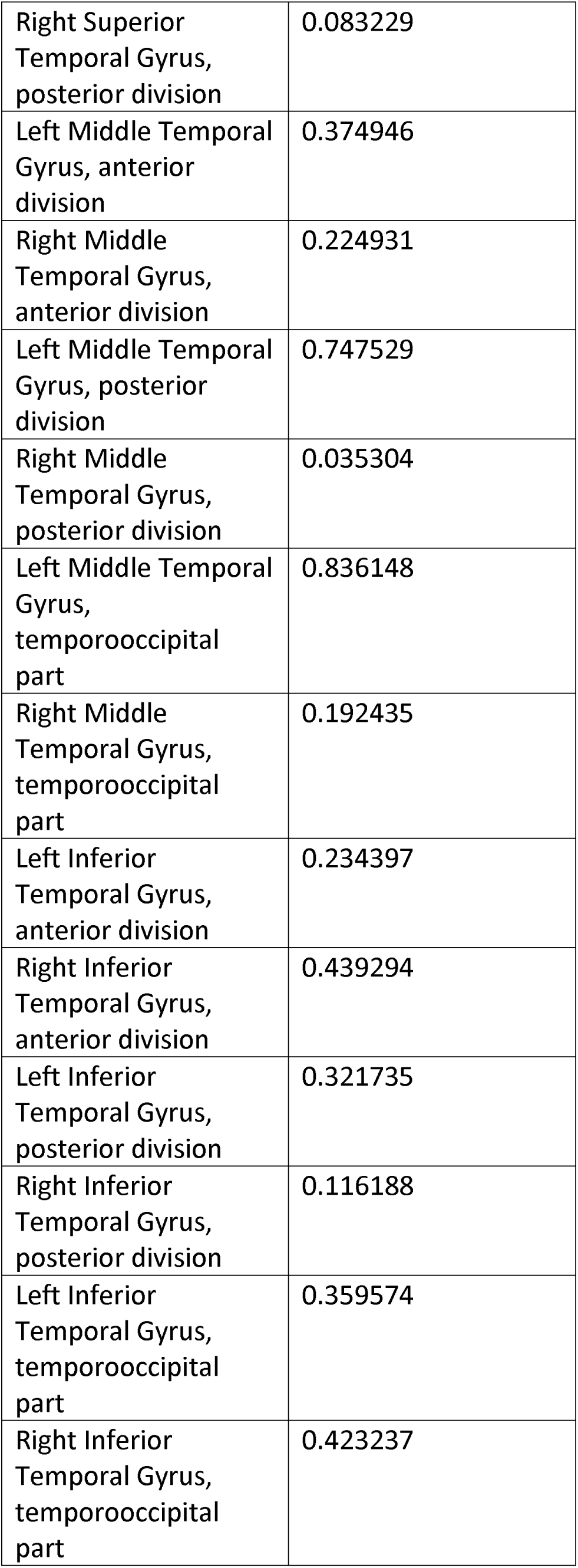
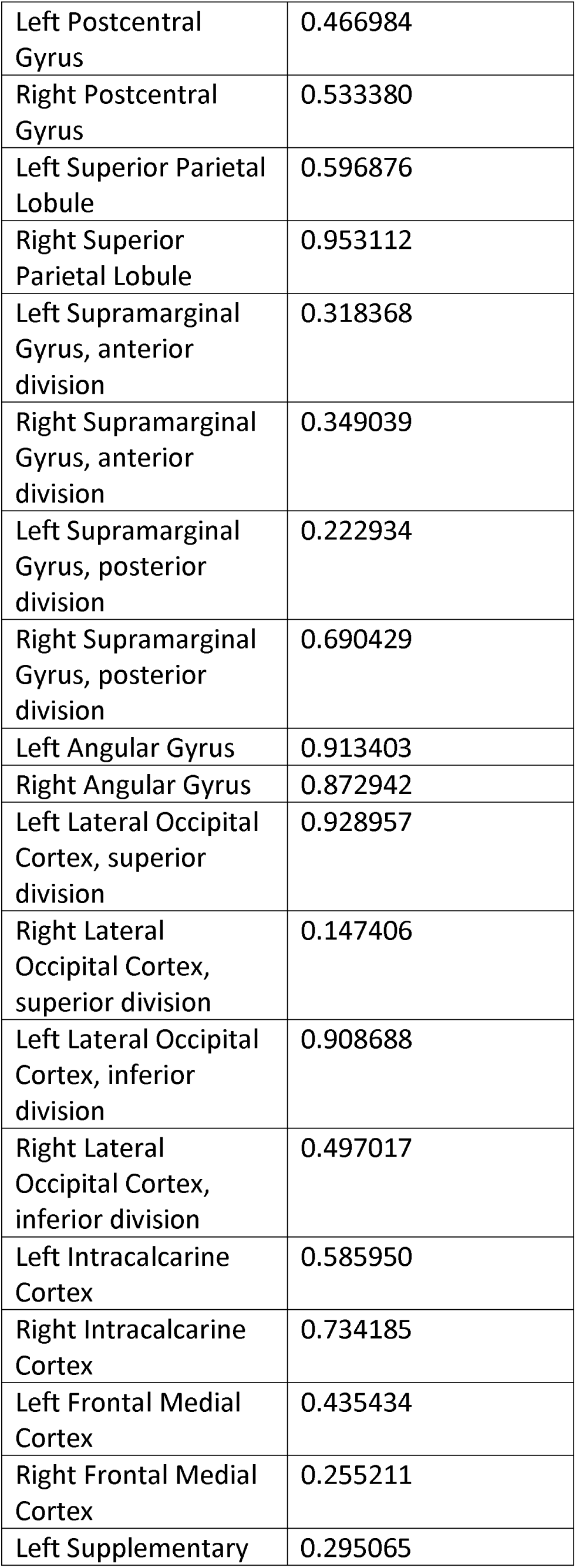
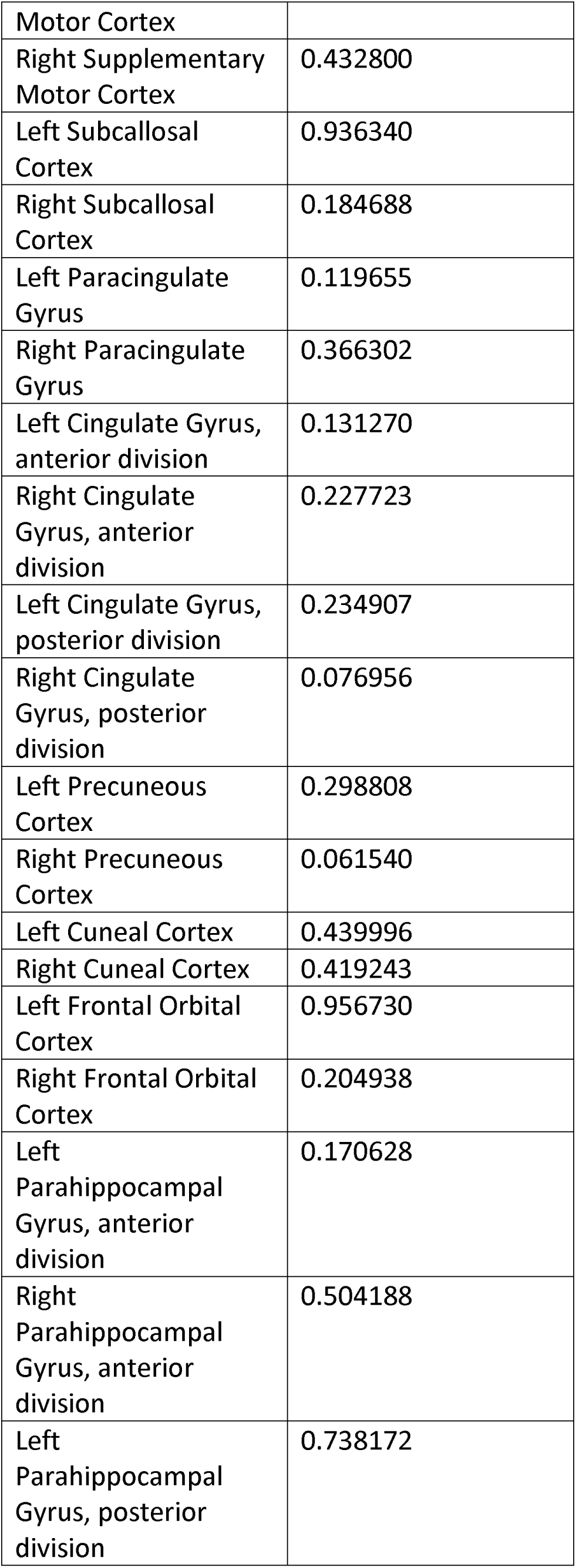
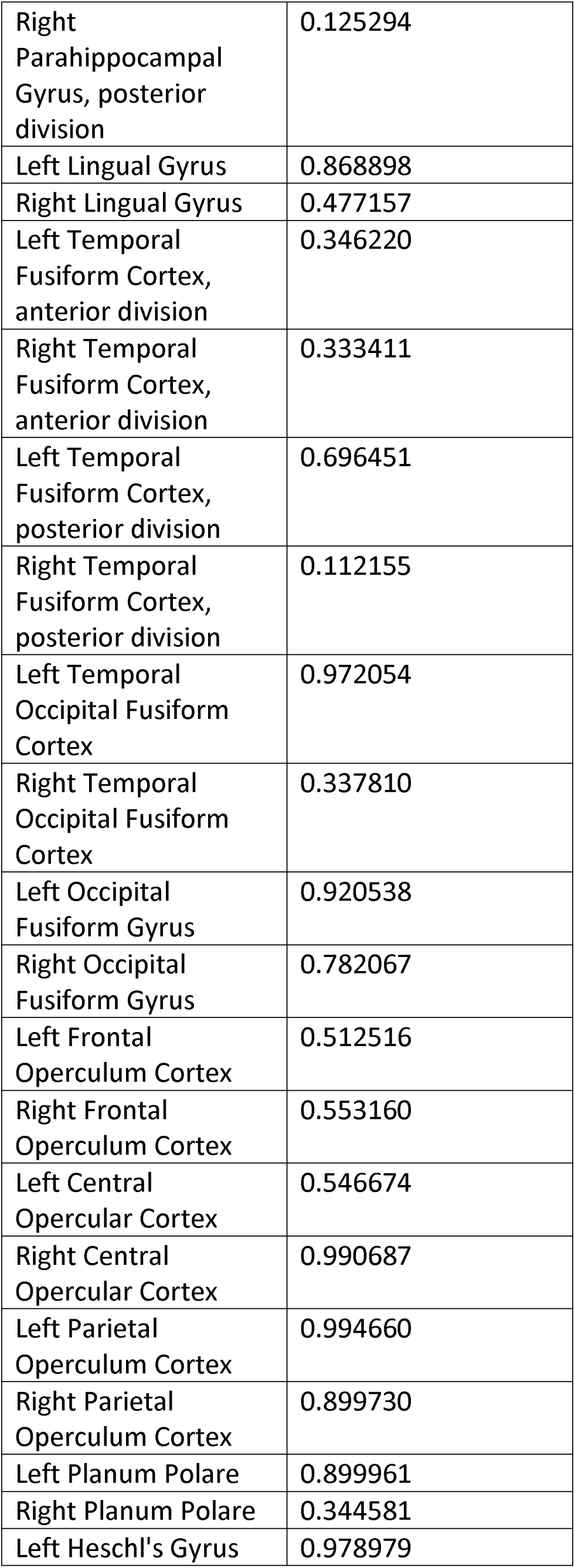
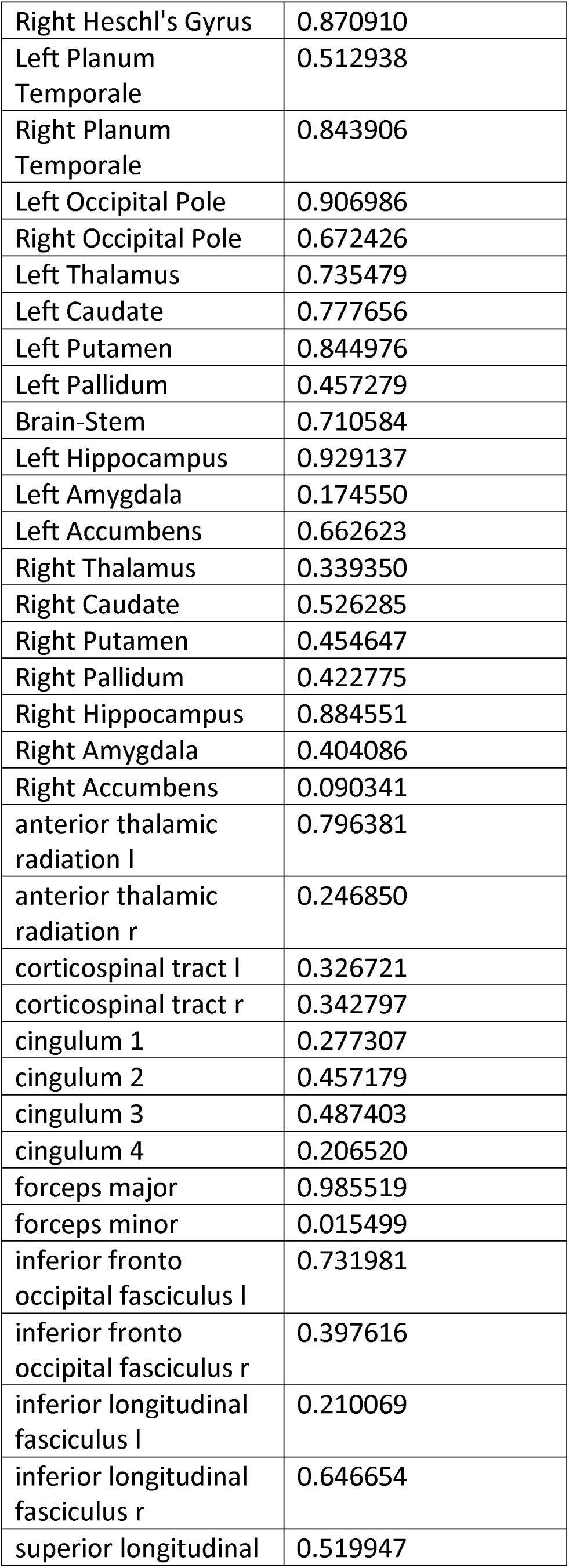
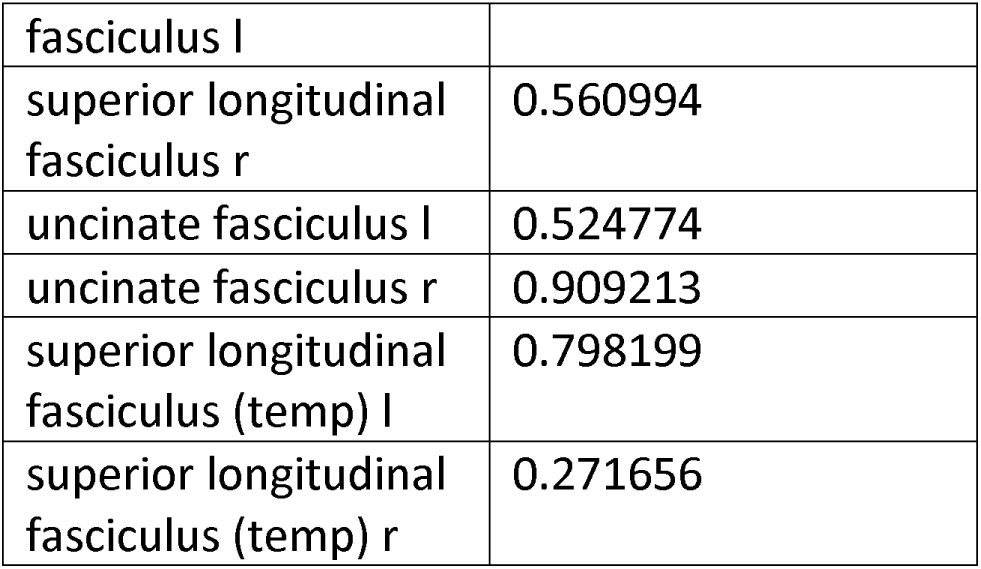
GASROS: Region-wise lesion load: Women versus Men.

**Supplementary Table 2.**
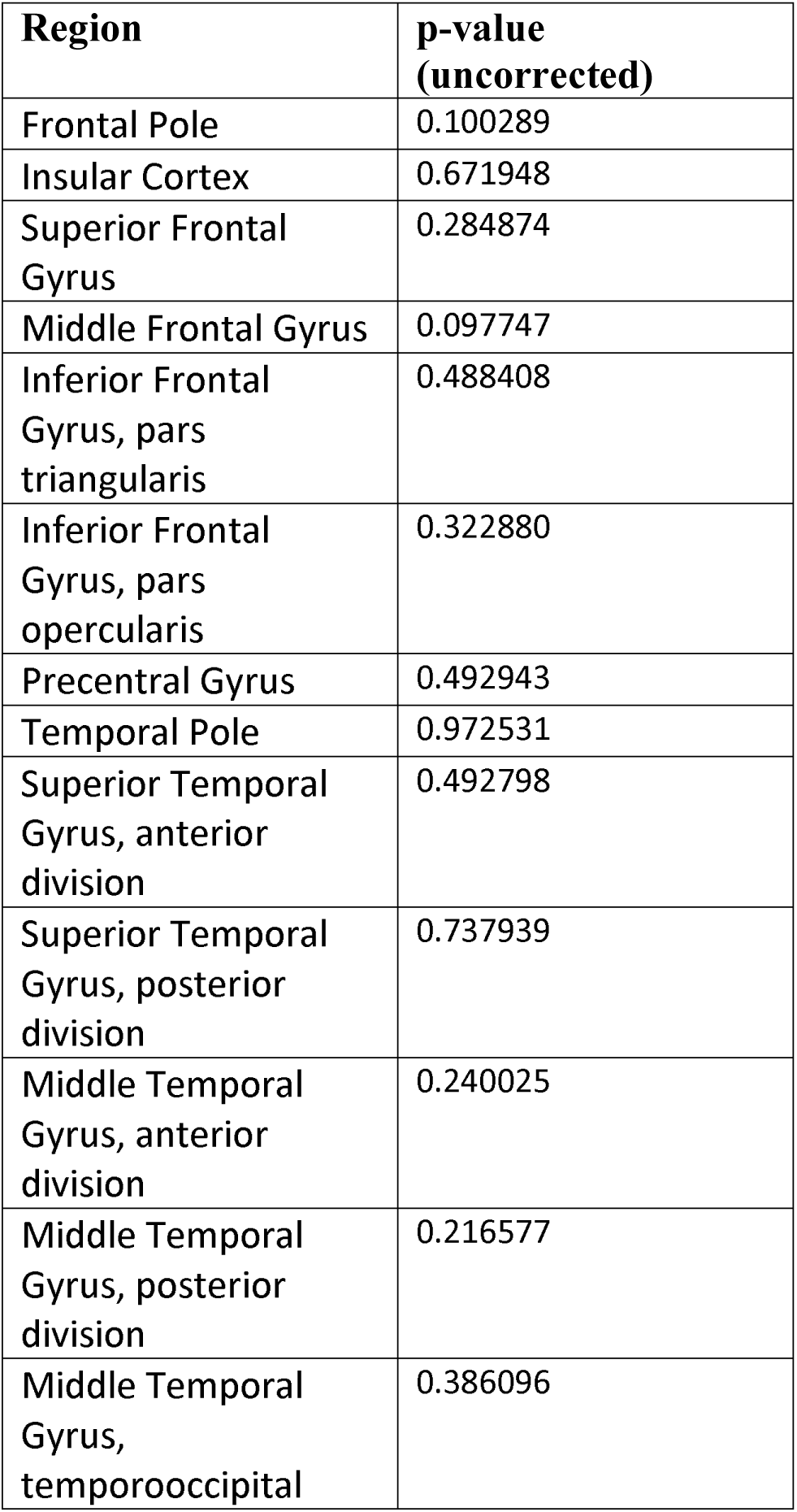
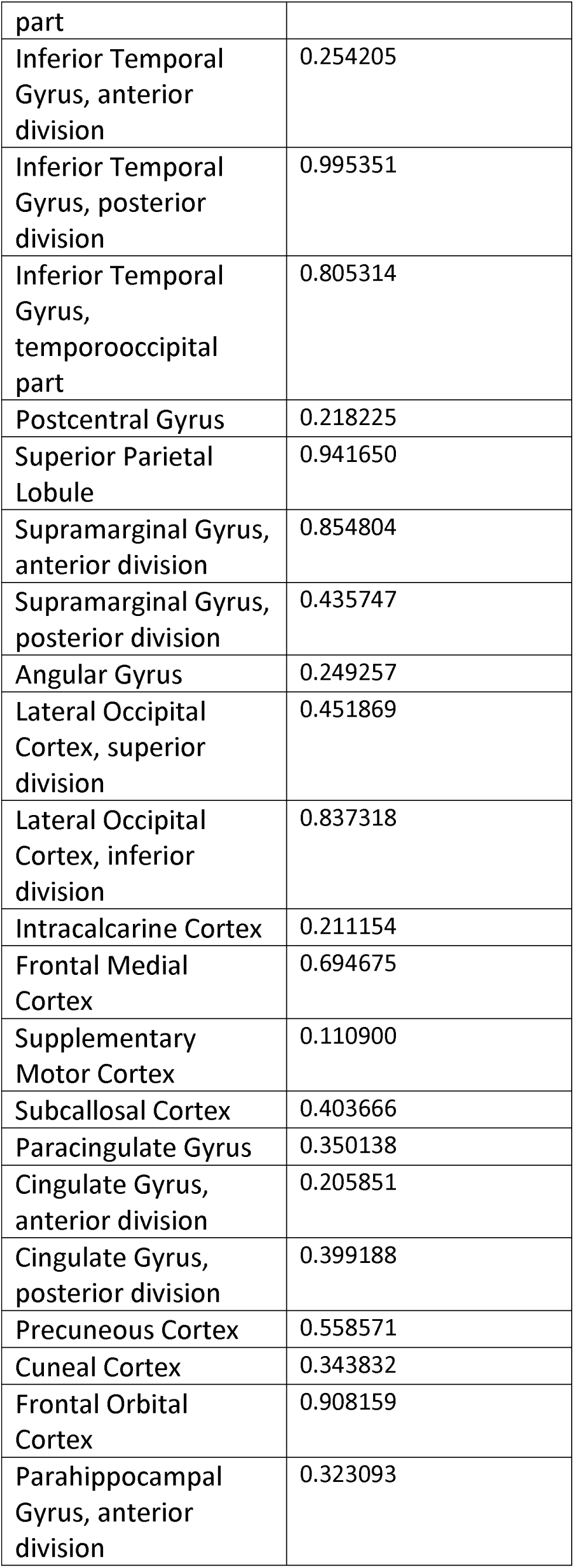
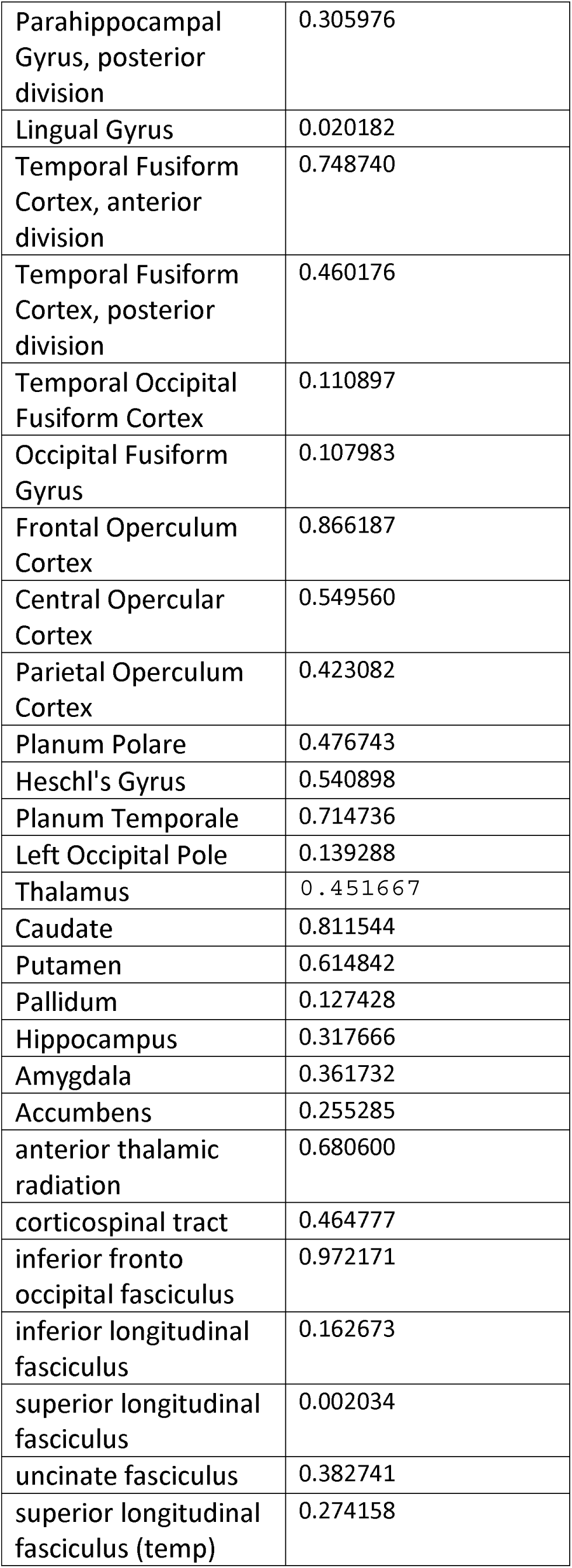
GASROS: Region-wise lesion load: Left versus Right hemisphere.

**Supplementary Table 3.** Matching of lesion atoms via correlations.

| Development cohort:<br>Number of lesion atom | Replication cohort: Number<br>of lesion atom | Pearson correlation of<br>NMF-weights: p-value |
| --- | --- | --- |
| 1 | 1 | $1.5e^{-26}$ |
| 2 | 1 and 4 | 1: $1.8e^{-8}$ and 2: $6.1e^{-12}$ |
| 3 | 3 | $1.4e^{-51}$ |
| 4 | 8 | $4.6e^{-10}$ |
| 5 | 4 | $1.8e^{-27}$ |
| 6 | 5 | $8.2e^{-45}$ |
| 7 | 6 (and 7) | 3: $1.9e^{-24}$ (and 9: $3.4e^{-11}$) |
| 8 | 9 | $2.1e^{-21}$ |
| 9 | 9 | $3.6e^{-19}$ |
| 10 | 10 | $1.5e^{-33}$ |

## References

1. Feigin, V. L. et al. Global and regional burden of stroke during 1990–2010: findings from the Global Burden of Disease Study 2010. The Lancet 383, 245–255 (2014).

2. Kelly-Hayes, M. et al. The influence of gender and age on disability following ischemic stroke: the Framingham study. Journal of Stroke and Cerebrovascular Diseases 12, 119–126 (2003).

3. Benjamin, E. J., et al. Heart disease and stroke statistics-2017 update: a report from the American Heart Association. Circulation 135, e146–e603 (2017).

4. Bushnell, C. et al. Sex differences in the evaluation and treatment of acute ischaemic stroke. The Lancet Neurology 17, 641–650 (2018).

5. Carcel, C., Woodward, M., Wang, X., Bushnell, C. & Sandset, E. C. Sex matters in stroke: A review of recent evidence on the differences between women and men. Frontiers in Neuroendocrinology 59, 100870 (2020).

6. Petrea, R. E. et al. Gender Differences in Stroke Incidence and Poststroke Disability in the Framingham Heart Study. Stroke 40, 1032–1037 (2009).

7. Bushnell, C. et al. Guidelines for the Prevention of Stroke in Women: A Statement for Healthcare Professionals From the American Heart Association/American Stroke Association. Stroke 45, 1545–1588 (2014).

8. Lisabeth, L. D., Brown, D. L., Hughes, R., Majersik, J. J. & Morgenstern, L. B. Acute stroke symptoms: comparing women and men. Stroke 40, 2031–2036 (2009).

9. Jerath, N. U., Reddy, C., Freeman, W. D., Jerath, A. U. & Brown, R. D. Gender differences in presenting signs and symptoms of acute ischemic stroke: a population-based study. Gender Medicine 8, 312–319 (2011).

10. Mandelzweig, L., Goldbourt, U., Boyko, V. & Tanne, D. Perceptual, Social, and Behavioral Factors Associated With Delays in Seeking Medical Care in Patients With Symptoms of Acute Stroke. Stroke 37, 1248–1253 (2006).

11. Beal, C. C. Women’s interpretation of and cognitive and behavioral responses to the symptoms of acute ischemic stroke. Journal of Neuroscience Nursing 46, 256–266 (2014).

12. Friberg, L., Benson, L., Rosenqvist, M. & Lip, G. Y. H. Assessment of female sex as a risk factor in atrial fibrillation in Sweden: nationwide retrospective cohort study. BMJ 344, e3522–e3522 (2012).

13. Lang, C. et al. Do Women With Atrial Fibrillation Experience More Severe Strokes?: Results From the Austrian Stroke Unit Registry. Stroke 48, 778–780 (2017).

14. Dehlendorff, C., Andersen, K. K. & Olsen, T. S. Sex Disparities in Stroke: Women Have More Severe Strokes but Better Survival Than Men. Journal of the American Heart Association 4, (2015).

15. Silva, G. S. et al. Gender Differences in Outcomes after Ischemic Stroke: Role of Ischemic Lesion Volume and Intracranial Large-Artery Occlusion. Cerebrovascular Diseases 30, 470– 475 (2010).

16. Hier, D. B., Yoon, W. B., Mohr, J. P., Price, T. R. & Wolf, P. A. Gender and aphasia in the stroke data bank. Brain and Language 47, 155–167 (1994).

17. Bates, E., et al. Voxel-based lesion–symptom mapping. Nature neuroscience 6, 448 (2003). 18.

18. Karnath, H.-O. The Anatomy of Spatial Neglect based on Voxelwise Statistical Analysis: A Study of 140 Patients. Cerebral Cortex 14, 1164–1172 (2004).

19. Sperber, C. Rethinking causality and data complexity in brain lesion-behaviour inference and its implications for lesion-behaviour modelling. Cortex 126, 49–62 (2020).

20. Wu, O. et al. Role of acute lesion topography in initial ischemic stroke severity and long-term functional outcomes. Stroke 46, 2438–2444 (2015).

21. Bonkhoff, A. K., et al. Generative lesion pattern decomposition of cognitive impairment after stroke. bioRxiv (2020).

22. Lee, D. D. & Seung, H. S. Learning the parts of objects by non-negative matrix factorization. Nature 401, 788 (1999).

23. Renoux, C. et al. Confounding by Pre-Morbid Functional Status in Studies of Apparent Sex Differences in Severity and Outcome of Stroke. Stroke 48, 2731–2738 (2017).

24. Koellhoffer, E. C. & McCullough, L. D. The Effects of Estrogen in Ischemic Stroke. Translational Stroke Research 4, 390–401 (2013).

25. McKinlay, S. M., Brambilla, D. J. & Posner, J. G. The normal menopause transition. Maturitas 14, 103–115 (1992).

26. Giese, A.-K., et al. Design and rationale for examining neuroimaging genetics in ischemic stroke: The MRI-GENIE study. Neurology Genetics 3, e180 (2017).

27. Hausmann, M. Why sex hormones matter for neuroscience: A very short review on sex, sex hormones, and functional brain asymmetries. Journal of Neuroscience Research 95, 40–49 (2017).

28. Hirnstein, M., Hugdahl, K. & Hausmann, M. Cognitive sex differences and hemispheric asymmetry: A critical review of 40 years of research. Laterality: Asymmetries of Body, Brain and Cognition 24, 204–252 (2019).

29. McGlone, J. Sex differences in human brain asymmetry: A critical survey. Behavioral and brain sciences 3, 215–227 (1980).

30. Inglis, J., Ruckman, M., Lawson, J. S., MacLean, A. W. & Monga, T. N. Sex differences in the cognitive effects of unilateral brain damage. Cortex 18, 257–275 (1982).

31. Inglis, J. & Lawson, J. S. A meta-analysis of sex differences in the effects of unilateral brain damage on intelligence test results. Canadian Journal of Psychology/Revue canadienne de psychologie 36, 670 (1982).

32. Inglis, J., Ruckman, M., Lawson, J. S., MacLean, A. W. & Monga, T. N. Sex differences in the cognitive effects of unilateral brain damage: Comparison of stroke patients and normal control subjects. Cortex 19, 551–555 (1983).

33. Cheng, B. et al. Influence of Stroke Infarct Location on Functional Outcome Measured by the Modified Rankin Scale. Stroke 45, 1695–1702 (2014).

34. Smith, D. V., Clithero, J. A., Rorden, C. & Karnath, H.-O. Decoding the anatomical network of spatial attention. Proceedings of the National Academy of Sciences 110, 1518–1523 (2013).

35. Siegel, J. S. et al. Disruptions of network connectivity predict impairment in multiple behavioral domains after stroke. Proceedings of the National Academy of Sciences 113, E4367–E4376 (2016).

36. Reeves, M. J. et al. Sex differences in stroke: epidemiology, clinical presentation, medical care, and outcomes. The Lancet Neurology 7, 915–926 (2008).

37. McCarthy, M. M. & Arnold, A. P. Reframing sexual differentiation of the brain. Nature Neuroscience 14, 677–683 (2011).

38. Hurn, P. D. & Macrae, I. M. Estrogen as a Neuroprotectant in Stroke. J Cereb Blood Flow Metab 20, 631–652 (2000).

39. Viscoli, C. M. et al. A clinical trial of estrogen-replacement therapy after ischemic stroke. New England Journal of Medicine 345, 1243–1249 (2001).

40. Nudy, M., Chinchilli, V. M. & Foy, A. J. A systematic review and meta-regression analysis to examine the ‘timing hypothesis’ of hormone replacement therapy on mortality, coronary heart disease, and stroke. IJC Heart & Vasculature 22, 123–131 (2019).

41. Alkayed, N. J. et al. Gender-linked brain injury in experimental stroke. Stroke 29, 159–165 (1998).

42. McCullough, L. D., Alkayed, N. J., Traystman, R. J., Williams, M. J. & Hurn, P. D. Postischemic Estrogen Reduces Hypoperfusion and Secondary Ischemia After Experimental Stroke. Stroke 32, 796–802 (2001).

43. Liu, M., Dziennis, S., Hurn, P. D. & Alkayed, N. J. Mechanisms of gender-linked ischemic brain injury. Restor Neurol Neurosci 27, 163–179 (2009).

44. Gibson, C. L. Cerebral ischemic stroke: is gender important? J Cereb Blood Flow Metab 33, 1355–1361 (2013).

45. Hara, Y. et al. Synaptic correlates of memory and menopause in the hippocampal dentate gyrus in rhesus monkeys. Neurobiology of Aging 33, 421.e17–421.e28 (2012).

46. Jacobs, E. G. et al. Reorganization of Functional Networks in Verbal Working Memory Circuitry in Early Midlife: The Impact of Sex and Menopausal Status. Cerebral Cortex bhw127 (2016) doi:10.1093/cercor/bhw127.

47. Kent, D. M., Price, L. L., Ringleb, P., Hill, M. D. & Selker, H. P. Sex-Based Differences in Response to Recombinant Tissue Plasminogen Activator in Acute Ischemic Stroke: A Pooled Analysis of Randomized Clinical Trials. Stroke 36, 62–65 (2005).

48. Shobha, N., Sylaja, P. N. & Kapral, M. K. Differences in stroke outcome based on sex. 7 (2010).

49. Lee, S.-J., Heo, S. H., Ambrosius, W. T. & Bushnell, C. D. Factors mediating outcome after stroke: gender, thrombolysis, and their interaction. Translational stroke research 9, 267– 273 (2018).

50. Sheth, S. A. et al. Sex Differences in Outcome After Endovascular Stroke Therapy for Acute Ischemic Stroke. Stroke (2019) doi:10.1161/STROKEAHA.118.023867.

51. Albers, G. W. et al. Thrombectomy for Stroke at 6 to 16 Hours with Selection by Perfusion Imaging. New England Journal of Medicine 378, 708–718 (2018).

52. Hinman, J. D. et al. Principles of precision medicine in stroke. Journal of Neurology, Neurosurgery & Psychiatry 88, 54–61 (2017).

53. Mauvais-Jarvis, F. et al. Sex and gender: modifiers of health, disease, and medicine. Lancet 396, 565–582 (2020).

54. Liu, S., Seidlitz, J., Blumenthal, J. D., Clasen, L. S. & Raznahan, A. Integrative structural, functional, and transcriptomic analyses of sex-biased brain organization in humans. Proceedings of the National Academy of Sciences 201919091 (2020) doi:10.1073/pnas.1919091117.

55. Smith, S. M. et al. Enhanced Brain Imaging Genetics in UK Biobank. bioRxiv (2020). 56.

56. Mah, Y.-H., Husain, M., Rees, G. & Nachev, P. Human brain lesion-deficit inference remapped. Brain 137, 2522–2531 (2014).

57. Etherton, M. R. et al. Sex-specific differences in white matter microvascular integrity after ischaemic stroke. Stroke and Vascular Neurology 4, 198–205 (2019).

58. Schirmer, M. D. et al. Rich-Club Organization: An Important Determinant of Functional Outcome After Acute Ischemic Stroke. Front. Neurol. 10, (2019).

59. Schirmer, M. D. et al. Spatial Signature of White Matter Hyperintensities in Stroke Patients. Frontiers in neurology 10, 208–208 (2019).

60. Rorden, C. & Brett, M. Stereotaxic display of brain lesions. Behavioural neurology 12, 191– 200 (2000).

61. Brudfors, M., Balbastre, Y., Nachev, P. & Ashburner, J. A Tool for Super-Resolving Multimodal Clinical MRI. arXiv preprint arXiv:1909.01140 (2019).

62. Ashburner, J. & Friston, K. J. Unified segmentation. Neuroimage 26, 839–851 (2005).

63. Schirmer, M. D., et al. White matter hyperintensity quantification in large-scale clinical acute ischemic stroke cohorts–The MRI-GENIE study. NeuroImage: Clinical 101884 (2019).

64. Desikan, R. S. et al. An automated labeling system for subdividing the human cerebral cortex on MRI scans into gyral based regions of interest. Neuroimage 31, 968–980 (2006).

65. Mori, S., Wakana, S., Van Zijl, P. C. & Nagae-Poetscher, L. M. MRI atlas of human white matter. (Elsevier, 2005).

66. Gelman, A. & Hill, J. Data analysis using regression and multilevel/hierarchical models. (Cambridge university press, 2006).

67. Schwartz, M. F., Faseyitan, O., Kim, J. & Coslett, H. B. The dorsal stream contribution to phonological retrieval in object naming. Brain 135, 3799–3814 (2012).

68. Hoffman, M. D. & Gelman, A. The No-U-Turn sampler: adaptively setting path lengths in Hamiltonian Monte Carlo. Journal of Machine Learning Research 15, 1593–1623 (2014).

69. Abraham, A. et al. Machine learning for neuroimaging with scikit-learn. Frontiers in neuroinformatics 8, 14 (2014).

70. Salvatier, J., Wiecki, T. V. & Fonnesbeck, C. Probabilistic programming in Python using PyMC3. PeerJ Computer Science 2, e55 (2016).

## References

Giese A-K, Schirmer MD, Donahue KL, Cloonan L, Irie R, Winzeck S, et al. Design and rationale for examining neuroimaging genetics in ischemic stroke: The MRI-GENIE study. Neurology Genetics 2017; 3: e180.

Wu O. A multi-center investigation of the association of acute stroke severity and long-term outcome with acute stroke lesion topography. In: JOURNAL OF CEREBRAL BLOOD FLOW AND METABOLISM. SAGE PUBLICATIONS INC 2455 TELLER RD, THOUSAND OAKS, CA 91320 USA; 2019. p. 38–38

Wu O, Winzeck S, Giese A-K, Hancock BL, Etherton MR, Bouts MJ, et al. Big Data Approaches to Phenotyping Acute Ischemic Stroke Using Automated Lesion Segmentation of Multi-Center Magnetic Resonance Imaging Data. Stroke 2019: STROKEAHA. 119.025373.

